# STK32A is a dual-specificity AGC kinase with a preference for acidic substrates

**DOI:** 10.1101/2020.03.04.976555

**Authors:** Fiona J. Sorrell, Fabrizio Miranda, Kamal R. Abdul Azeez, Apirat Chaikuad, Arminja N. Kettenbach, Scott A. Gerber, Stefan Knapp, Ahmed A. Ahmed, Jonathan M. Elkins

## Abstract

The STK32 kinases are a small subfamily of three uncharacterised serine/threonine kinases from the AGC kinase family whose functional role is so far unknown. Here, we analyse the consensus peptide sequence for STK32A phosphorylation, showing that STK32A is directed towards acidic substrate sequences and exhibits dual-specificity for serine/threonine and tyrosine residues. A crystal structure of STK32A reveals an overall structure typical of the AGC protein kinase family but with significant and unique features including an altered binding mode of the hydrophobic motif to the N-terminal lobe of the kinase domain, and a novel alpha-helix in between the turn motif and the hydrophobic motif. The crystal structure combined with phylogenetic analysis reveals the evolutionary conservation of the acidic substrate preference. *In vitro* binding assays demonstrated that the STK32 kinases bind significant numbers of clinically used kinase inhibitors.

## Introduction

The protein kinases STK32A, STK32B and STK32C, also known as YANK1, YANK2 and YANK3 (<u>Y</u>et <u>A</u>nother <u>N</u>ovel <u>K</u>inase), are adjacent to the well-studied Aurora kinases on the kinase phylogenetic tree [1] but by contrast very little is known about their functions. Currently the available published data largely comprises genetic associations, cellular localisation studies, and mouse knockout data. STK32A RNA is highly expressed in brain and endocrine tissues, while STK32A protein is localized to the centrosome [2]. STK32B is highly expressed in kidney and localized to microtubules, while STK32C protein is ubiquitously expressed [2]. STK32A knockout mice show increased monocytes and circulating LDL cholesterol, while STK32C knockout mice show abnormal locomotor behaviour [3].

Despite their poor functional annotation, single nucleotide polymorphisms (SNPs) point to important physiological functions of these kinases. For instance, polymorphisms in STK32A were linked to coeliac disease [4,5]. A methylation site within the STK32A gene was linked to smoking [6], while in a genome-wide association study (GWAS) the PPP2R2B-STK32A-DPYSL3 locus was found to be a susceptibility locus for lung cancer [7]. In an RNAseq study identifying genes regulated by the Wnt3a/β-catenin pathway, STK32A was slightly upregulated (1.8x) in response to Wnt3 treatment in hippocampal neurons [8]. In one study STK32B protein levels were found to be upregulated in more aggressive breast cancers [9], while another study found them to be downregulated in oral squamous cell carcinoma [10]. Driver mutations in various cancers in both STK32A and STK32B have been identified [11], including a G35E missense mutation on the glycine-rich phosphate-binding loop of STK32B detected in melanoma and an S89F missense mutation of STK32A, also in melanoma. Of significant interest as the only study looking at chemical inhibition of these kinases, by examining the effects of a large number of compounds on the THP-1 cell line, STK32C was identified as a potential anti-target for small molecule kinase inhibitor development, due to a possible relationship with cytotoxicity [12].

STK32B and STK32C have both been linked in genetic studies to several neurological disorders. In a genome-wide association study (GWAS) for essential tremor in a European population a single nucleotide polymorphism (SNP) in STK32B was associated with essential tremor [13], an observation replicated in a Chinese population [14]. The European study also demonstrated increased expression of STK32B in the cerebellar cortex in essential tremor patients [13]. Some patients with Ellis-van Creveld syndrome have a deletion of an entire 520-kb region including STK32B in addition to EVC and EVC2; it is unknown to what, if any, extent the loss of STK32B contributes to the syndrome [15]. In the same genomic region of chromosome 4p16, an association between SNPs in STK32B and EVC with on-syndromic oral clefts was observed [16]. In a study of monozygotic twins a differentially methylated position in STK32C was found to be associated with adolescent depression [17]. An SNP within STK32C was also linked to risk of psychiatric disorders in Down Syndrome patients [18].

As well as being largely unstudied, the human STK32s are not close in sequence to other human kinases. For example, STK32A has 70% and 67% sequence identity over the kinase domain to STK32B and STK32C respectively, but only 36% sequence identity to the next closest homologues (PRKACG or RSK2). To initiate the functional characterisation of these proteins we determined the X-ray crystal structure of STK32A and a solution-phase envelope using small angle X-ray scattering (SAXS), identified a consensus substrate motif for STK32A, and analysed the binding of a set of kinase inhibitors, including some that have been used in the clinic.

## Results

### Conservation and evolution of STK32A, STK32B and STK32C

Sequence analysis identified homologues of STK32A in fungi, including *Magnaporthe oryzae* (rice blast fungus), *Neurospora crassa* (red bread mould) and *Schizosaccharomyces pombe* (fission yeast), indicating it is conserved across opisthokonts (Figure 1A). STK32B likely evolved more recently, with homologues indicated in bilateral animals including nematodes (*Caenorhabditis elegans*), jawless vertebrates (such as *Eptatretus burgeri* (hagfish) and *Petromyzon marinus* (Lamprey)) and some insects (for example *Drosophila melanogaster* (fruit fly) and *Acyrthosiphon pisum* (pea aphid)). STK32C appears to be the newest gene, present only in bony vertebrates. The classification of Manning et al. placed the STK32s adjacent to the Aurora kinases on the phylogenetic tree [1], but unlike Aurora A/B/C the STK32 kinases are genuine AGC kinases, having the full C-terminal extension typical of the AGC family. This C-terminal region contains the hydrophobic motif (ϕ-X-X-ϕ-S/T-ϕ, ϕ=hydrophobic), an important regulatory element known to stabilise the active conformation of AGC kinases. The serine or threonine residue within the canonical hydrophobic motif is a site of activating phosphorylation in most AGC kinases, however in the STK32 family this site corresponds to an asparagine in vertebrates and in some bilateral animals such as *Drosophila*, or an aspartate in fungi (Figure 1B). N-terminal to the hydrophobic motif is the turn motif which is well-conserved from humans to nematodes (Figure 1B). In fungi, however, the potential phosphorylatable serine residue in the turn motif (highlighted in bold in Figure 1B) is exchanged for an acidic glutamate residue, which may mimic the phosphorylated residue, and in fission yeast (*S.p.* PPK33) the entire C-terminal extension is truncated prior to the turn motif. Between the turn motif and the hydrophobic motif secondary structure prediction indicates the presence of a long α-helical element that is unique to the STK32s and well conserved across the entire family from fungi to humans (residues 346-370 in human STK32A), which here we have termed the “HF motif helix” (Figure 1B).

**Figure 1.**
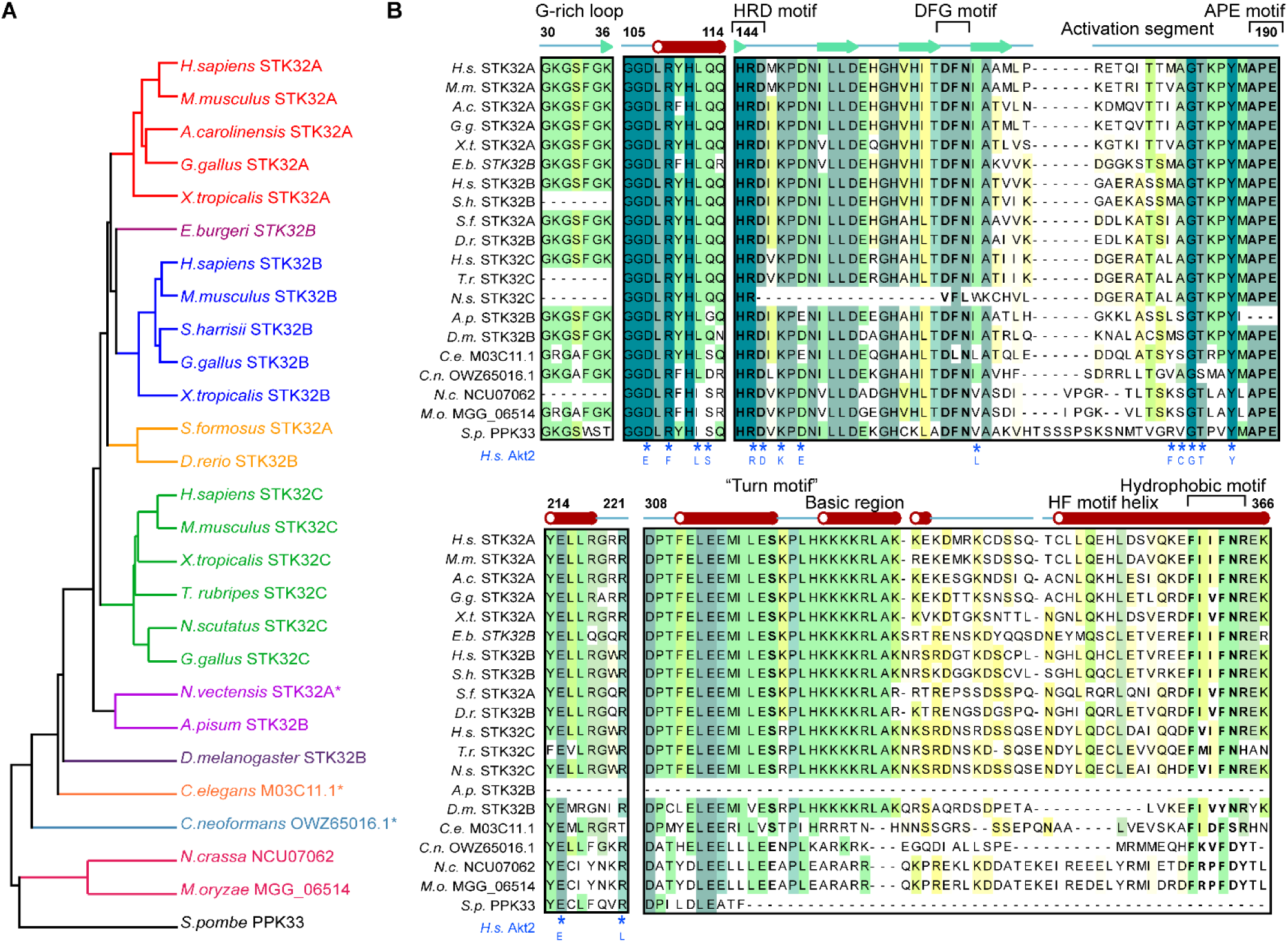
Evolution of STK32A, STK32B and STK32C and homologues. **(A)** Phylogenetic tree for STK32 homologues in human (*Homo sapiens, H.s.*), mouse (*Mus musculus, M.m.*), anole lizard (*Anolis carolinensis, A.c.*), red junglefowl (*Gallus gallus, G.g.*), Western clawed frog (*Xenopus tropicalis, X.t.*), hagfish (*Eptatretus burgeri, E.b.*), Tasmanian devil (*Sarcophilus harrisii, S.h.*), Asian bony tongue (*Schleropages formosus, S.c.*), zebrafish *(Danio rerio, D.r.*), fugu (*Takifugu rubripes, T.r.*), mainland tiger snake (*Notechis scutatus, N.s.*), starlet sea anemone (*Nematostella vectensis, N.v.*), pea aphid (*Acyrthosiphon pisum, A.p.*), fruit fly (*Drosophila melanogaster, D.m.*), nematode (*Caenorhabditis elegans, C.e.*), encapsulated yeast (*Cryptococcus neoformans, C.n*), red bread mould (*Neurospora crassa, N.c.*), rice blast fungus (*Magnaporthe oryzae, M.o*) and fission yeast (*Schizosaccharomyces pombe*). * indicates putative or uncharacterised protein sequence. **(B)** Sequence alignment of STK32A homologues across multiple organisms and comparison with STK32B and STK32C. Predicted secondary structural elements for human STK32A are indicated (red cylinder = α-helix; green arrow = β-sheet; blue line = unstructured loop). Important residues and conserved motifs are highlighted in bold. Blue stars indicate residues involved in substrate binding in *H.s.* Akt2 mapped onto the sequence, with the Akt2 equivalent residue shown underneath. Sequences are coloured by conservation where darker colour indicates higher conservation.

As well as these AGC kinase-specific motifs, the STK32 kinases contain the typical sequence motifs of active protein kinases, such as a glycine-rich phosphate-binding loop (“G-rich loop”), the HRD (His-Arg-Asp) motif vital for phosphate transfer, and the APE (Ala-Pro-Glu) motif that secures the C-terminal end of the activation segment. The DFG (Asp-Phe-Gly) motif, required for proper positioning of ATP for phosphate transfer, is also present in the STK32 family, although the glycine is substituted for a less conformationally flexible asparagine residue which is remarkably conserved. While red bread mould (*N.c.* NCU07062) shows relatively high sequence conservation to the STK32 family across the C-terminal half of the kinase domain, the turn motif and hydrophobic motif, the N-terminus is truncated so that the αC helix and glycine-rich phosphate-binding loop are completely absent suggesting that kinase activity may have been lost. The glycine-rich loop is also absent from some vertebrates, for example Tasmanian devil (*S. harrisii* STK32A/B/C), fugu (*T. rubripes* STK32C) and mainland tiger snake (*N. scutatus* STK32A/C), which otherwise share high similarity (60-80%) with the canonical human STK32 sequences. Human STK32A shares 69% and 65% sequence identity with STK32B and STK32C, respectively, and the key motifs are highly conserved. A notable difference is the presence of a 67-residue N-terminal extension in STK32C isoform 1 that is rich in proline, alanine, arginine and serine, therefore likely to be highly flexible, which is not present in any human STK32A or STK32B isoform. Several of the serine residues in this extension are predicted to be phospho-sites. Several other isoforms of both human STK32B and STK32C have been described with N-terminal truncations that remove the glycine-rich loop, similar to the homologues described above, and it may be that these isoforms are incapable of ATP binding and catalysis.

One feature that appears to be specific to STK32 kinases over the rest of the human kinome is a highly basic region immediately following the turn motif. In vertebrates this region (residues 324-333 in human STK32A) consists of the conserved sequence HKKKKRLA(K/R)(X/-)(K/R), whereas in more primitive organisms this region is less tightly conserved but generally contains a higher proportion of arginine and fewer lysine residues (Figure 1B). The function of this basic patch is currently unknown.

### STK32A is an active dual-specificity Ser/Thr and Tyr kinase that preferentially phosphorylates acidic substrates

Most AGC kinases have been reported to preferentially phosphorylate substrates containing basic residues such as arginine or lysine N-terminal to the serine/threonine phosphorylation site, and even share multiple common substrates [19]. Several crystal structures of AGC kinases in complex with their substrate peptides exist, for example AKT2 in complex with a GSK3 peptide motif [20], therefore it is possible to predict the residues in STK32A that may be involved in substrate binding through sequence homology (Figure 1B, blue stars). The residues in the AKT2 substrate binding cleft that directly contact the substrate (annotated in blue in Figure 1B) appear to be relatively well conserved in STK32A/B/C and therefore based on the sequence alone it would have been a reasonable hypothesis that the STK32 family also preferentially phosphorylates basic substrates, similar to other AGC kinases.

STK32A kinase domain protein was expressed in insect cells using baculoviruses. Expression yields were good and the protein was efficiently purified by nickel-affinity chromatography followed by size exclusion chromatography (SEC). The protein eluted from the SEC column at the expected volume for a monomer. The resultant protein was auto-phosphorylated following expression, but lambda phosphatase treatment enabled the production of unphosphorylated protein for use in assays and structural studies (Figure 2A).

**Figure 2.**
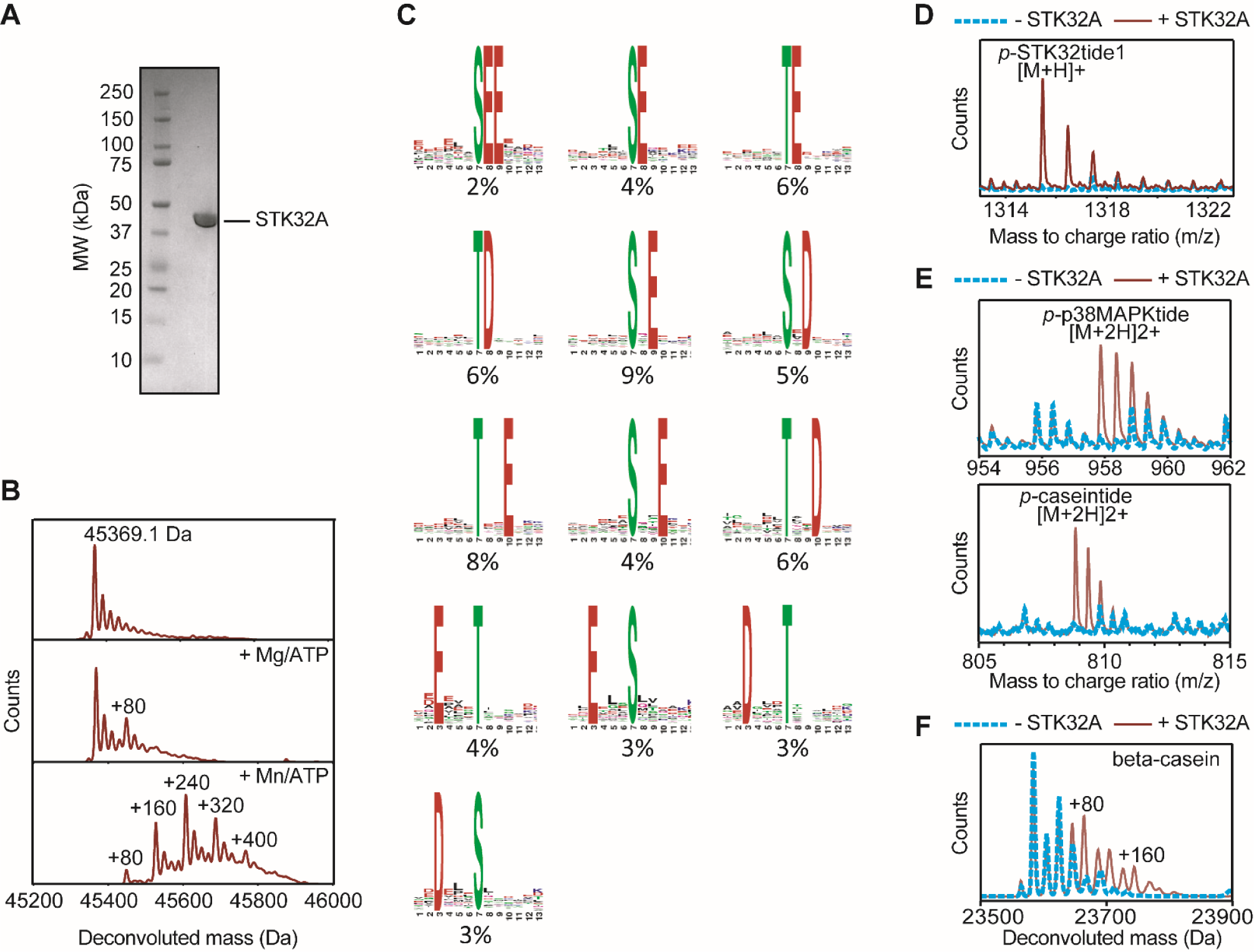
STK32A activity, substrate recognition screening and peptide phosphorylation. **(A)** SDS-PAGE analysis of purified STK32A protein typical of that used for subsequent studies. **(B)** Auto-phosphorylation of STK32A after 1 hour incubation with ATP and magnesium (centre) or manganese (bottom), compared to control (top), measured by LC-MS. **(C)** Representative panels of STK32A substrate specificity defining peptide residues identified using in vivo isolation. Each panel represents a separate clustering of peptide sequences, with the most commonly observed residues at each position at the top of each letter stack. The percentage of peptides in each cluster are indicated below each panel. **(D)** LC-MS analysis of synthetic consensus peptide substrate STK32tide1 after phosphorylation by STK32A (red solid line) compared to control without STK32A (blue dotted line) **(E)** LC-MS analysis of synthetic peptide substrates corresponding to activation loop sequence of p38-MAPK (top) and bovine beta-casein (bottom) after phosphorylation by STK32A (red solid line) compared to control without STK32A (blue dotted line). **(F)** LC-MS of full-length bovine beta-casein incubated in the absence of STK32A (blue dotted line) and following phosphorylation by STK32A (red solid line).

The ability of purified, dephosphorylated, STK32A kinase domain to auto-phosphorylate *in vitro* was assessed by mass spectrometry. Auto-phosphorylation occurred more rapidly in the presence of manganese than magnesium; after one hour incubation with 5 mM ATP, STK32A was auto-phosphorylated on up to five residues in the presence of Mn^2+^, compared to a single site in the presence of Mg^2+^ (Figure 2B). The sites of phosphorylation were mapped following digestion of the protein using elastase which indicated that STK32A auto-phosphorylates at multiple sites: S33, Y49, T163, S227, T229, S230, S231, S320, S354 and T383 (Supplementary Figure 1). These sites include phosphorylation on the typical AGC kinase turn motif site (S320) and close to the hydrophobic motif at S354. A cluster of serine and threonine residues predicted to be situated on the αF-αG loop, a structural element usually important for interaction with substrates, was also found to be auto-phosphorylated *in vitro* (S227, T229, S230, S231).

We incubated purified STK32A kinase domain with peptides derived from proteolysed and stringently dephosphorylated HeLa cell lysates and analysed the resulting phosphorylation patterns by mass spectrometry [21]. This revealed a set of consensus substrate recognition sequences for STK32A (Figure 2C). The data revealed that STK32A phosphorylates serine or threonine residues, and has a general preference for acidic residues (aspartate or glutamate) at position P-4, P+1, P+2 and P+3, and no other significant preferences.

Based on these consensus substrate recognition motifs, two synthetic peptides were designed to test the ability of STK32A to phosphorylate a specific substrate with an acidic residue in positions P-4, P+1, P+2 and P+3. These peptides were designated “STK32tide-1, corresponding to the amino acid sequence ASEALV<u>S</u>EEDAD, and “STK32tide-2”, corresponding to ADELL<u>S</u>EVEAKK. After incubation of 250 µM of each peptide for 18 hours with STK32A and ATP, species indicative of the phosphorylated STK32tide-1/2 sequences were observed by LC-MS (Figure 2D and Supplementary Figure 1A). Given that phosphorylated products were only detectable after a relatively long incubation time, the phosphorylation of these peptides by STK32A appears to be quite inefficient, but nevertheless confirms that STK32A is able to phosphorylate *in vitro* even peptides with multiple acidic residues across positions -4 and +1 to +3. Similar to the observation that auto-phosphorylation proceeds faster in the presence of manganese than magnesium, the peptides were also phosphorylated more rapidly in the presence of manganese (data not shown), substantiating a general preference for manganese as the counter-ion.

Next, a small in-house library of kinase substrate peptides was tested against STK32A at a concentration of 500 µM. CDC25Ctide, designed to correspond to residues 193-205 of M-phase inducer phosphatase 3 (also known as dual specificity phosphatase CDC25C), covering the known phosphorylation site of PLK1 and PLK3 (ELMEF<u>S</u>LKDQEAK), was found to be weakly phosphorylated by STK32A after a long incubation time (Supplementary Figure 2A), similar to the designed STK32tide peptides, and contains an acidic residue in the P+3 position. Caseintide, a synthetic peptide corresponding to the acidic sequence KKEKFQ<u>S</u>EEQQQ and designed to mimic part of bovine beta-casein, was phosphorylated by STK32A after only 2 hours, and contains acidic residues in positions P-4, P+1 and P+2 (Figure 2E, bottom). Also after 2 hours a species corresponding to phosphorylated p38MAPKtide was detected (Figure 2E, top). The sequence of p38MAPKtide corresponds to part of the activation loop of p38α (MAPK14) (AGAGLARH<u>T</u>DDEM<u>T</u>G<u>Y</u>VA) and contains three potential phosphorylation sites (corresponding to p38α residues T175, T180 and Y182). Based on the consensus sequence data we hypothesized that the two most likely STK32A phosphorylation sites in this peptide corresponded to T175 (containing acidic residues in positions P+1, P+2 and P+3) and T180 (containing an acidic residue in position P-4). Interestingly, however, mass spectrometry analysis revealed that sites T180 and Y182 were phosphorylated by STK32A, and there was no evidence of T175 phosphorylation (Supplementary Figure 2B and Supplementary Table 1). Several peptides were detected that contained dual phosphorylation both at the T180 and Y182 sites. This indicated that STK32A is a dual specificity serine/threonine and tyrosine kinase.

After seeing the ability of STK32A to phosphorylate both caseintide and the p38α synthetic peptides, we tested whether STK32A was also capable of phosphorylating the corresponding full-length proteins *in vitro*. A sample of dephosphorylated casein extracted from bovine milk, containing an approximate ratio of 5:3:1 of beta-casein to alpha-S1-casein to kappa-casein, was purchased. After 2 hours incubation of the casein extract in the presence of STK32A, ATP and a mixture of manganese/magnesium chloride, up to two additions of 80 Da mass to beta-casein was evident, indicating STK32A was capable of phosphorylating full-length beta-casein (Figure 2F). Conversely, STK32A did not appear to be capable of phosphorylating full-length p38 *in vitro*. Recombinantly expressed p38α (MAPK14) was mixed with STK32A, ATP and either manganese chloride or a mixture of manganese and magnesium chloride, both in the presence and absence of the specific p38α inhibitor Skepinone-L. After 24 hours in the absence of Skepinone-L, there was evidence of phosphorylation of p38α, but not when Skepinone-L was added, indicating that the observed activity was likely due to auto-phosphorylation by p38α itself rather than STK32A (Supplementary Figure 2C). STK32A was capable of auto-phosphorylation in the presence of Skepinone-L, confirming that Skepinone-L did not affect the activity of STK32A (Supplementary Figure 2D). Therefore, although STK32A was able to phosphorylate the synthetic peptide derived from the p38α activation loop, full-length p38α did not appear to be a substrate for STK32A *in vitro*.

### The crystal structure of human STK32A demonstrates canonical AGC-kinase as well as STK32-specific structural features

Crystallisation trials were conducted using STK32A (residues P9-D390) pre-mixed with the non-specific kinase inhibitor staurosporine. It was necessary to use a relatively high STK32A concentration of 27 mg/mL (0.6 mM) to obtain optimal crystals. STK32A:staurosporine crystallised in space group *P*3_2_ with six STK32A molecules in the asymmetric unit, each bound to one molecule of staurosporine. Each of the six molecules in the asymmetric unit superimposed well, with minor differences around the flexible C-terminal extension turn motif (described in more detail below). Data collection and refinement statistics are shown in Table 1. In the final model, refined to a resolution of 2.29 Å, residues 15-373 were resolved in the electron density, except for a flexible loop containing the basic patch (residues 323-332) which was disordered.

**Table 1.** Data collection and refinement statistics.

|  |  |
| --- | --- |
|  | STK32A :<br>Staurosporine |
| PDB ID | 4FR4 |
| Space group | $P3_2$ |
| No. of molecules in the asymmetric unit | 6 |
| Unit cell dimensions<br>$a, b, c$ (Å), $\alpha, \beta, \gamma$ (°) | 154.1, 154.1, 112.1<br>90.0, 90.0, 120.0 |
| **Data collection** |  |
| Resolution range (Å)<sup>a</sup> | 51.68-2.29<br>(2.41-2.29) |
| Unique observations<sup>a</sup> | 128336 (19231) |
| Average multiplicity<sup>a</sup> | 3.3 (3.2) |
| Completeness (%)<sup>a</sup> | 95.7 (97.8) |
| $R_{\text{merge}}$<sup>a</sup> | 0.09 (0.64) |
| Mean $\langle(I)/\sigma(I)\rangle$<sup>a</sup> | 11.2 (2.0) |
| **Refinement** |  |
| Resolution range (Å) | 50.44-2.29 |
| $R$-value, $R_{\text{free}}$ | 0.21, 0.24 |
| r.m.s. deviation from ideal bond length (Å) | 0.015 |
| r.m.s. deviation from ideal bond angle (°) | 1.48 |
| Ramachandran Outliers<sup>b</sup> | 0.05% |
| Most favoured | 94.06% |

<sup>a</sup> Values within parentheses refer to the highest resolution shell.
<sup>b</sup> Values from Molprobit.

The model revealed that STK32A shares canonical structural features of protein kinases, with an N-terminal lobe consisting of five beta-strands and two alpha-helices, and a mostly helical C-terminal lobe. The ATP binding site, occupied by the inhibitor staurosporine is between the N-lobe and the C-lobe (Figure 3A). STK32A also retains the canonical feature of AGC kinases, a C-terminal extension that wraps around the N-terminal kinase lobe, secured by the interaction of the hydrophobic motif (HF motif) with the groove next to the αC-helix, stabilising the αC-helix in an active conformation (Figure 3B).

**Figure 3.**
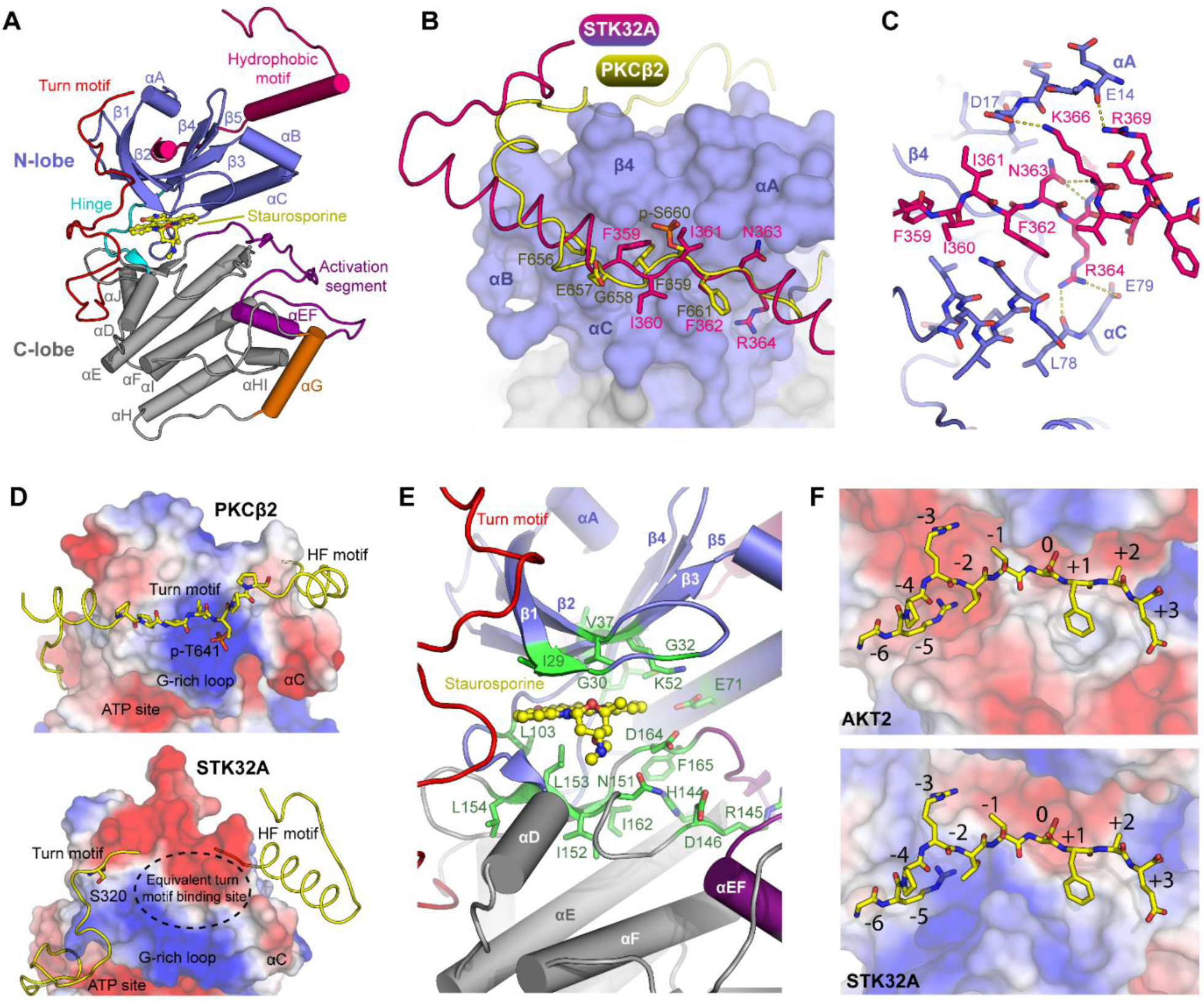
STK32A has an AGC kinase fold with several unique features. **(A)** Overview of STK32A structure showing typical features of AGC kinases as highlighted. **(B)** Hydrophobic (HF) motif binding in STK32A (pink) differs from other typical AGC kinases such as PKCβ2 (superposed in yellow) with a ‘register shift’ of two residues, putting PKCβ2 pSer660 in the same binding position as STK32A Ile361 rather than the equivalent residue Asn363, as well as changes in backbone orientation. **(C)** Detailed view of the interactions between the HF motif (pink) and N-lobe (blue) in STK32A with key hydrogen bonds shown by yellow dotted line. **(D)** Comparison of the turn motif binding site between PKCβ2 and STK32A showing differences in electrostatic potential across the surface (blue = positive charge, red = negative charge, white = neutral). **(E)** Detailed view of ATP-binding site in STK32A showing highly conserved residues important for ATP-binding and catalysis in green. STK32A was co-crystallised with the broad spectrum kinase inhibitor Staurosporine in the ATP-binding site, shown in yellow. **(F)** Comparison of the substrate binding groove of STK32A with AKT2: (top) Crystal structure of AKT2 bound to peptide substrate (yellow) (PDB 1O6K) and (bottom) superposition of the peptide from 1O6K onto the STK32A structure where several steric clashes and repulsive charge interactions are evident. Vacuum electrostatics were generated using Pymol.

The canonical AGC kinase HF motif is ϕ-X-X-ϕ-S/T-ϕ (ϕ = hydrophobic residue, S/T = serine or threonine phosphorylation site) however STK32A/B/C have F-X-X-F-N-R (Figure 1B). The lack of the canonical phosphorylation site and the presence of the HF bound to the N-lobe showed that unlike some other AGC kinases STK32A/B/C do not require an activating phosphorylation on the C-terminal tail to form an active conformation. During the activation process of many AGC kinases such as SGKs, S6Ks, RSKs or PKCs, the phosphorylated hydrophobic motif binds to the PIF pocket for PDK1, promoting phosphorylation of the activation loop by PDK1 [19]; it is currently unknown whether PDK1 is involved in the activation of STK32A/B/C.

The binding mode of the HF motif to the N-lobe of STK32A is significantly different to that of other AGC kinases. Compared to the binding of example canonical HF motifs such as PKCβ2 or PKCι the binding positions of the key hydrophobic interactions are altered. In PKCβ2 F656 is the first hydrophobic residue of the HF motif and slots into a conserved pocket on the αC helix to help position this important element into the active conformation (Figure 3B). The first HF motif hydrophobic residue in STK32A is F359, which however binds in the equivalent location to PKCβ2 F659, the second hydrophobic residue of the HF motif. Phosphorylated S660 in PKCβ2 points away from the kinase N-lobe (Figure 3B) and in STK32A this position is occupied by I361 (Figure 3B). Situated just after the HF motif are a series of residues that form polar interactions with αA and αC to secure the C-terminal region across the top of the N-lobe. STK32A K366 is well conserved across the STK32 family and forms a hydrogen bond to D17 from the N-terminus. Nearby, R369 interacts with the backbone carbonyl of E14 and R364 forms a hydrogen bond with the backbone of L78 and with the side chain of E79, further securing this region in position (Figure 3C).

Two conformations of the C-terminal tail around the turn motif are evident in the crystal structure: half of the molecules in the asymmetric unit (Chains D,E,F) favour one conformation where residues P309-S320 form an ordered α-helix that packs against the N-lobe β-sheet formed of strands β1/β2/β3, and the other molecules (Chains A,B,C) form a loop that packs against the αC and HF-motif helices and activation loop of a nearby symmetry-related monomer (Supplementary Figure 3A). Part of the basic turn motif, comprising multiple lysine residues, is unresolved in the electron density (a.a. 323-333) in all chains, indicating the high flexibility of this region. Other typical AGC kinases such as PKCβ2 (PDB 2I0E) and PKCι (PDB 1ZRZ) contain a positively charged binding pocket for the phosphorylated turn motif, normally required for full kinase activity [22], but this appears to be lacking in STK32A, which instead contains acidic or neutral residues at the equivalent site (Figure 3D). It is possible that STK32s do not contain a true turn motif, although we have mapped one auto-phosphorylation site of STK32A to S320 at the usual turn motif site (Supplementary Figure 1). It seems plausible that the phosphorylated turn motif binds in an alternative binding pocket in STK32s, for example the basic patch immediately on top of the glycine-rich loop, adjacent to the normal turn motif binding site in other AGC kinases. There is a possibility that the basic sequence (HKKKKRLAKK) in STK32s immediately following the traditional turn motif (Figure 1) interacts with the acidic patch observed on the N-lobe (Figure 3D) as a further means of securing the C-terminus in position, however this region was disordered in the crystal structure and, therefore, appeared to be highly flexible. Upon turn motif phosphorylation in STK32s it is possible that this region would adopt a more ordered conformation.

At the ATP binding site, the relatively small gatekeeper residue (V100) affords larger than average capacity at the back of the binding pocket, potentially explaining the ability of STK32A to bind inhibitors designed for gatekeeper mutant kinases (see inhibitor binding analysis below). Otherwise STK32A maintains the typical set of hydrophobic residues in β1 and β2 of the N-lobe required to bind the adenosine moiety of ATP (I29, G30, V37 and A50), as well as the highly conserved residues at the hinge and C-lobe (L103, I152, L153, L154) (Figure 3E) that form the base of the adenosine binding pocket in the structures of most active protein kinases [23]. Highly conserved polar residues that stabilise the γ-phosphate of ATP and divalent cation are also conserved in STK32A (K52, E71, D146, K148, N151, D164) (Figure 3E). Other conserved residues found throughout the structural core of active-conformation kinases, including the AGC kinases AKT2 and PKA [23] are also retained in STK32A, although larger variation is evident compared to AKT2/PKA (Supplementary Table 3).

Interestingly, the key sites that have been shown to be responsible for the interaction of AKT2 with peptide substrates seem to be conserved in STK32A (Figure 1), at first glance indicating that STK32A should be capable of interacting with basic substrates similar to AKT2 and other AGC kinases, however inspection of the residues surrounding the substrate binding groove in the crystal structure indicates a potential explanation for the observed preference for acidic substrates: situated within close proximity to the substrate binding groove are basic residues Arg109, Arg221 and Arg304. These sites are occupied by acidic or hydrophobic residues in AKT2, allowing the arginine side chains of the substrate to form salt bridges or participate in pi-pi stacking (Figure 3F, top panel). If this basic peptide substrate is modelled into the substrate binding groove of STK32A the side-chains of Arg109 and Arg304 superimposed closely with the position of the side-chains of the arginine residues in positions -3 and -5 of the peptide substrate (Figure 3F, bottom panel), indicating that binding of this type of substrate to STK32A in a similar orientation may have unfavourable charge-charge interactions. Modelling of the acidic STK32tide-1 peptide into the crystal structure (Supplementary Figure 3B) indicated favourable interactions at the substrate binding groove and offered a structural hypothesis as to why STK32A is able to phosphorylate this substrate *in vitro*. The substrate binding grooves of STK32A, STK32B and STK32C are highly conserved (Supplementary Figure 3C) and therefore all may bind acidic substrates in a similar manner to STK32A.

### Solution phase analysis of STK32A

We performed a Small Angle X-ray Scattering (SAXS) analysis of the same unphosphorylated STK32A protein that was crystallised to assess the solution phase behaviour of the protein, and also compared this to the auto-phosphorylated protein (Figure 4, Supplementary Table 3). The proteins were passed through a size-exclusion chromatography column immediately before SAXS analysis. The proteins eluted as a single, sharp peak with consistent radius of gyration (*R*_*g*_) calculated across the peak for both unphosphorylated and phosphorylated protein (Figure 4A, 4B). In general, the solution scattering behaviour of both samples was very similar as seen in the similar scattering curves, Guinier plots and overlapping real space p(r) distributions of each (Figure 4C-E). A plateau in the Porod-Debye plot (Figure 4F) and a Porod exponent approaching 4 indicated that the proteins were relatively compact and rigid, as expected for a globular protein kinase domain. The normalised Kratky plots for both samples had a peak maximum approaching that expected by Guinier’s approximation (√3 with magnitude 1.04) and returning to baseline at higher values of *q* x *R*_*g*_, indicating that they are generally globular and folded, but the slight shift of the peak to the right indicated that they are slightly elongated (Figure 4G). Measurements revealed that the protein was monomeric in solution, both in its unphosphorylated form and following auto-phosphorylation. A mass of 47-51 kDa was determined by SAXS by the *Q*_*R*_ method [24] for the unphosphorylated protein, compared to an intact mass of 45.4 kDa observed by denaturing LC-MS (Supplementary Table 3). A small mass increase was detectable by SAXS following auto-phosphorylation, consistent with addition of multiple phosphate groups, as observed by LC-MS (Supplementary Table 3). The experimental SAXS data was compared to calculated SAXS curves based on the STK32A crystal structure which had missing loops modelled (Model 1), and resulted in an excellent fit (Chi^2^ = 1.18), indicating that the conformation observed for each inhibitor-bound monomer in the crystal structure was not very dissimilar to the conformation in solution (Figure 4H). *Ab initio* dummy residue models generated using the SAXS data could also be superimposed well with the envelope of the crystal structure (Figure 4I, 4J). The *ab initio* model calculated for the phosphorylated protein sample retained a generally similar overall shape to that of the unphosphorylated protein (Figure 4K). Altogether, the data indicates that STK32A functions as a monomer in both its unphosphorylated and auto-phosphorylated state.

**Figure 4.**
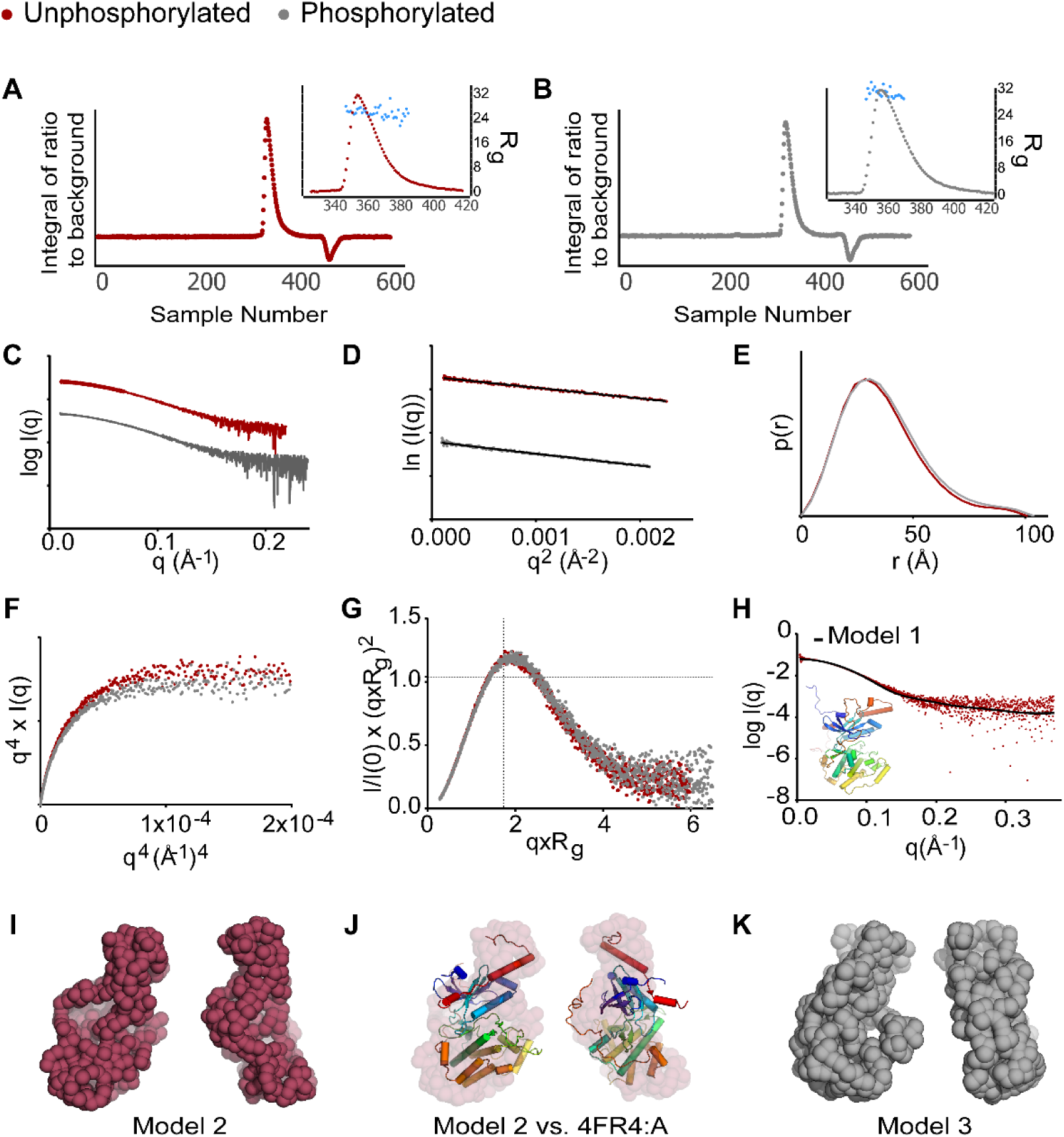
Solution phase analysis of unphosphorylated (dark red) and auto-phosphorylated (grey) STK32A. **(A)** HPLC-SAXS data for unphosphorylated STK32A and (inset) radius of gyration versus sample number across the eluted peak. **(B)** Equivalent of (A) for the auto-phosphorylated protein. **(C)** SAXS q versus log I(q) scattering profile for the major peak of phosphorylated versus unphosphorylated STK32A, offset for clarity. **(D)** Guinier/reciprocal space plots of both samples, offset for clarity. **(E)** Normalised p(r) distribution. **(F)** Porod-Debye plot. **(G)** Normalised Kratky plot. **(H)** Fit of Model 1 versus raw SAXS data for unphosphorylated STK32A. **(I)** *Ab initio* dummy residue model generated for unphosphorylated STK32A. **(J)** Superposition of crystallographic model versus SAXS dummy residue model. **(K)** *Ab initio* dummy residue model generated for phosphorylated STK32A.

### STK32A binds multiple clinically-used kinase inhibitors

Differential scanning fluorimetry (DSF) measurements were used to assess inhibitor binding to STK32A by measuring the increase in melting temperature (ΔT_m_) of STK32A in the presence of a set of kinase inhibitors many of which have been used in clinical trials. STK32A bound several broad-spectrum type-I kinase inhibitors such as Staurosporine, TAE684, AZD7762 and K252a, but also many clinically-tested type-I inhibitors including the ALK inhibitor Ceritinib (LDK378), the BRAF inhibitor Dabrafenib (GSK2118436), the PAK4 inhibitor PF-03758309, the SYK inhibitor PRT062607, the p38 MAPK inhibitor TAK 715, and the Aurora A/B/C inhibitor Danusertib (Figure 5A). Interestingly, the compound 1NM-PP1, developed to target “analogue-sensitive” kinases [25] such as v-Src-as1 that harbours an I338G gatekeeper mutation, bound STK32A with ΔT_m_ = 8°C, the third highest thermal stabilisation observed with this set of inhibitors (Figure 5B). STK32A also bound several other inhibitors designed to target analogue sensitive kinases, such as PP-121, 2,3-dMB-PP1 and 1-Na-PP1 (Figure 5B). Several crystal structures exist for human and parasitic kinases in complex with similar small molecules that shed light on the likely binding mode of 1NM-PP1 in STK32A (for example 4LGH, 5W8R, 3I7B, 3NCG, 3MA6). In a crystal structure of analogue sensitive Src (PDB 4LGH), the pyrazolo-pyrimidine core binds to the kinase hinge and the naphthalene group extends into the buried back pocket of the ATP site. In STK32A, there is a small gatekeeper residue (Val100), conserved across the family (Supplementary Figure 3C), that leaves a small pocket at the back of the ATP-binding site that is unoccupied in the structure with Staurosporine (Figure 5C, circled), but potentially allows it to accommodate the bulky naphthalene group of 1NM-PP1 (Figure 5D). Out of 420 kinase sequences analysed, most human kinases contain a large or branched hydrophobic residue as the gatekeeper (Met (40%), Leu (17%), Phe (15%)), whereas only 22% contain a small residue (such as Thr, Ser, Val, Ala, Gly) and out of these valine was found to be the gatekeeper in only 3% of cases, predominantly consisting of tyrosine kinases, therefore the STK32 family is relatively unusual in this respect [26]. A similar binding mode is observed for PP-121 in Src (PDB 3EN4), however PP-121 is also able to form additional polar interactions between the nitrogen atoms of the pyrrolo-pyridine substituent and the side chain of Glu310 and the backbone of Asp404 in the back pocket of Src, residues that are conserved in STK32A (Glu71 and Asp164) further explaining the ability of STK32A to bind PP-121 relatively strongly (ΔT_m_ = 6°C).

**Figure 5.**
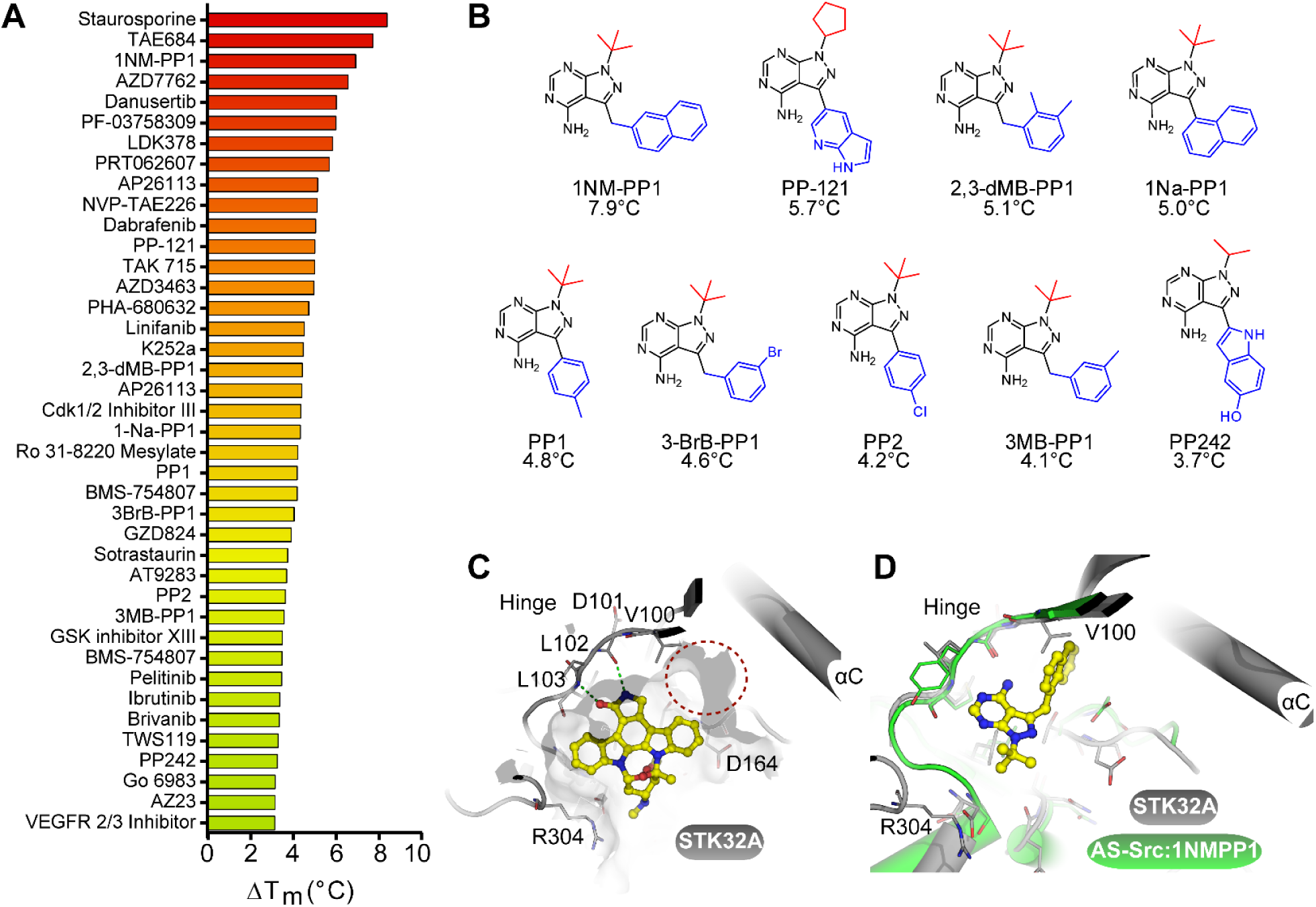
STK32A binds various clinically used kinase inhibitors as well as inhibitors designed to bind kinases with a small gatekeeper residue. **(A)** Inhibitors were assessed using a thermal shift assay and those that raised the melting temperature (T_m_) of STK32A by more than 3.5 °C are shown. **(B)** Structures of 1NM-PP1 (top left), an inhibitor that typically only binds to kinases having a small gatekeeper residue (Val100 in STK32A), and related compounds. The changes in T_m_ upon incubation with STK32A are listed underneath. **(C)** View of the STK32A ATP-binding site with Staurosporine (yellow) bound at the kinase hinge. Polar interactions are indicated with green dotted lines. A red circle indicates space at the back of the ATP binding site near the gatekeeper residue Val100 that potentially allows for binding of 1NM-PP1. **(D)** Superposition of the STK32A crystal structure (grey) with that of analogue-sensitive Src (“AS-Src”, shown in green, PDB 4LGH) bound to 1NM-PP1 (yellow).

## Discussion

The data shows that STK32A is an acidophilic, dual-specificity AGC-family kinase. Serine/threonine kinases that have been found to have a preference for acidic substrates are relatively rare, but include the casein kinases (CKs) and polo-like kinases (PLKs), all of which are able to use casein as a substrate. Similarly STK32A is able to phosphorylate casein in vitro; both our designed synthetic substrate corresponding to part of the bovine beta-casein peptide sequence as well as a sample of beta-casein extracted from bovine milk were phosphorylated by STK32A. Inspection of crystal structures of acidophilic kinases sheds light on their preference for negatively charged residues surrounding the substrate phosphorylation site, for example CK2α is known to phosphorylate a large number of substrates with acidic residues C-terminal to the phosphorylation site [27] and examination of crystal structures of CK2α reveals a large patch of highly positive charge on the surface close to the ATP-binding site that drives this preference (Supplementary Figure 4A). Similar patches of positive charge can be seen in the PLKs and other casein kinases as well as in our structure of STK32A (Supplementary Figure 4A). In CK2, a phosphorylated residue can substitute for acidic glutamate or aspartate residues in the substrate. The tyrosine kinase epidermal growth factor receptor (EGFR), which has also been found to be acidophilic in substrate preference in the P-3 to P-1 region, is similarly capable of binding primed, phosphorylated substrates [27,28]. EGFR contains a large basic patch across the substrate binding groove that could complement the binding of peptides with acidic residues N-terminal to the phosphorylation site, and a crystal structure solved with a primed phosphorylated substrate shows that the large phosphorylated tyrosine in the P+1 position is able to also occupy this pocket in a binding mode distinct from non-primed substrates (Supplementary Figure 4B) [29]. The phospho-mapping data for STK32A indicates that it may also be capable of binding primed substrates since there were found to be doubly-phosphorylated peptides situated both on auto-phosphorylated STK32A (S228/T229) and on the p38 synthetic peptide (residues equivalent to T180/Y182 in full-length p38), with sites within one or two amino acids of each other (Supplementary Figures 1 and 2).

STK32B and STK32C have 69% and 65% sequence identity, respectively, to STK32A. In particular, the residues lining the substrate binding groove of STK32A are highly conserved in STK32B/C, including Arg109, Arg221 and Arg304. Therefore, the general preference of STK32A for acidic substrates is expected to be conserved in STK32B/C. However, the αG helix and the central portion of the activation segment, also important for substrate recognition, are not well conserved (Supplementary Figure 3C) and there will be a range of different substrates for the three proteins. The expression patterns and subcellular localizations of STK32A, STK32B and STK32C will significantly affect the available substrates. Data in the Human Protein Atlas suggests that STK32A is located to the centrosome, while STK32B is mainly localised to microtubules and vesicles [30], while preliminary localisation studies show that with high level ectopic expression the STK32 proteins remain primarily cytosolic (Supplementary Figures 5 and 6).

The ATP-binding site is well conserved within the STK32 family (Supplementary Figure 3C), therefore a similar pattern of inhibitor binding is likely. In particular, the small valine gatekeeper residue is conserved and therefore inhibitors such as 1NM-PP1 and PP-121 (Figure 5B) are also expected to bind to STK32B and STK32C as well as STK32A. This may offer a convenient route to obtaining selective inhibitors of the STK32 family. We also observed that many well-known kinase inhibitors bind STK32A. Given that there is currently no information on the mechanistic roles of the STK32 kinases in the cell it is unclear if any of the observable phenotypes with these inhibitors are due in part to STK32 inhibition.

## Methods

### Cloning

DNA for residues 9-390 of human STK32A isoform 1 (NCBI reference NP_001106195.1) was PCR amplified and subcloned into baculovirus transfer vector pFB-LIC-Bse by ligation independent cloning. The resulting construct expressed the desired protein with an N-terminal hexahistidine purification tag and TEV (tobacco etch virus) protease tag cleavage site (extension MGHHHHHHSSGVDLGTENLYFQ*SM where * represents the TEV protease digestion site). The construct was verified by DNA sequencing.

### Protein Expression and Purification

The expression construct was transformed into *E. coli* DH10Bac competent cells, which were used to prepare recombinant bacmid DNA. The bacmid DNA was transfected into *Spodoptera frugiperda* insect cells (*Sf9)* from which recombinant baculovirus was recovered. After three cycles of amplification of the virus, *Sf9* cells at 2 million cells per mL were infected with virus for 48 hours at 27 °C. The cells were harvested by centrifugation at 1000 x g, re-suspended in Binding Buffer (50 mM Tris·HCl pH 7.8, 200 mM NaCl, 20 mM imidazole, 0.5 mM TCEP and protease inhibitor cocktail (Sigma)) and frozen at −30 °C until further use.

The re-suspended cells were thawed, lysed by sonication on ice, polyethylenimine (PEI) was added to a final concentration of 0.05%, and the insoluble debris was removed by centrifugation. The supernatant was passed through a column of 6 mL Ni-Sepharose resin (GE Healthcare) at 4 °C. The resin was washed with 100 mL Binding Buffer, 50 mL Binding Buffer containing 40 mM imidazole and 1 M NaCl, and 40 mL of Binding Buffer containing 60 mM imidazole, before elution with Binding Buffer containing 250 mM imidazole. TEV protease, MnCl_2_ (1 mM) and lambda phosphatase were added to the eluate and the final wash fraction, which were left overnight at 4 °C. The eluate and final wash fractions were combined, concentrated to 5 mL volume, and injected onto an S200 16/60 gel filtration column (GE Healthcare) pre-equilibrated into GF Buffer (20 mM Tris·HCl pH 7.8, 150 mM NaCl, 0.5 mM TCEP) at a flow rate of 1.0 mL/min. Fractions containing STK32A were further purified by passing through a gravity column of 0.8 mL Ni-Sepharose. The flow-through was collected and the column was further eluted with GF Buffer containing 10, 20, and 30 mM imidazole. Protein identities were confirmed by electrospray ionization mass spectrometry (ESI-MS), expected 45101.0 Da, observed 45101.9 Da.

### Consensus substrate identification

Peptides derived from HeLa cell lysate digests that were phosphorylated by STK32A were identified as in published procedures [21].

### *In vitro* phosphorylation reactions

STK32A auto-phosphorylation was tested by incubation of unphosphorylated purified protein at 2 mg/mL (34 µM) for 1 hour in the presence of 5 mM ATP and 5 mM cation (either magnesium chloride or manganese chloride) in buffer A (50 mM HEPES pH 7.5, 150 mM NaCl, 5% (w/v) glycerol, 0.5 mM TCEP and 100 µM sodium orthovanadate). Each reaction was quenched by 30 x dilution into 0.1% formic acid and loaded onto an electrospray ionisation time-of-flight (LC-MSD TOF) spectrometer (Agilent).

Two peptides based on the consensus substrate sequences for STK32A were synthesised: “STK32tide-1” (ASEALV<u>S</u>EEDAD) and “STK32tide-2” (ADELL<u>S</u>EVEAKK), as well as a small in-house synthetic library based on known common kinase substrates. The ability of STK32A to phosphorylate the peptides was tested by incubation in buffer A, supplemented with 5 mM MgCl_2_ or 5 mM MnCl_2_ and 5 mM ATP. The STK32tide peptides were tested at a final concentration of 250 µM, whilst the library peptides were tested at 500 µM. Reactions were quenched after 1, 2 or 18 hours following initiation of the reaction, and peptide phosphorylation was monitored by LC-MS as before.

For testing the phosphorylation of full-length p38α by STK32A, p38α (MAPK14) was recombinantly co-expressed in *E. coli* with lambda phosphatase. The purified p38α was mixed with STK32A and 5 mM MnCl_2_ or 5 mM MnCl_2_/MgCl_2_ and 5 mM ATP in buffer A, in the presence of either the specific p38α inhibitor Skepinone-L at 50 µM or 2.5% DMSO as a control. Reactions were quenched and monitored by LC-MS as before. For testing phosphorylation of casein by STK32A, a sample of dephosphorylated lyophilised bovine casein (Sigma-Aldrich product number C4032) was purchased and diluted to a final assay concentration of 1 mg/mL in buffer A supplemented with 5 mM MgCl_2_/MnCl_2_/ATP and 1.8 µM STK32A. Results were analysed by LC-MS as before.

### Crystallisation and data collection

The fractions containing pure STK32A were pooled, and staurosporine was added. The sample was concentrated to 36 mg/mL, diluted to 5 mL volume with GF Buffer and then re-concentrated to 36 mg/mL. This sample was then mixed 3:1 with peptide SEALVSEEDAD (at 34.4 mM in 80 mM Tris·HCl pH7.8) to give a final STK32A concentration of 27 mg/mL and a final peptide concentration of 8.6 mM. This sample was used for crystallisation. Note that although included in the crystallisation, the peptide was not visible in the electron density for the structure.

Crystals were obtained using the sitting drop vapour diffusion method at 4 °C. Crystals of grew from a mixture of 50 nL STK32A:staurosporine and 100 nL of a well solution containing 0.2M sodium acetate, 20% PEG 3350 and 10% ethylene glycol. Crystals were equilibrated into reservoir solution plus 25% ethylene glycol before freezing in liquid nitrogen. The data was collected at 100K at the Diamond Synchrotron, beamline I04-1. Data collection statistics can be found in Table 1. Protein concentrations were measured by UV absorbance, using the calculated molecular weights and estimated extinction coefficients using a NanoDrop spectrophotometer.

### Structure determination

The diffraction data was indexed and integrated using MOSFLM [31] and scaled using SCALA [32]. The structure was solved by molecular replacement using PHASER [33] and an ensemble search model created using PHENIX [34] from the kinase domain structures of human PKCiota (PDB 3A8X), cAMP-dependent protein kinase catalytic subunit alpha (PDB 3L9N) and p90 ribosomal S6 kinase 2 (PDB 3G51). There were six molecules of YANK1 in the asymmetric unit. The model was built using Coot [35] and refined with REFMAC5 [36] for 300 cycles of jellybody refinement prior to further refinement in PHENIX [34]. Refinement statistics are given in Table 1. Coordinates were deposited in the PDB under accession code 4FR4.

### HPLC-SAXS

Sample details, structural parameters and modelling results are given in Supplementary Table 1. Purified, unphosphorylated, STK32A (as used for the crystal structure determination) was concentrated to 10.5 mg/mL (∼230 µM) and flash frozen in liquid nitrogen. For preparation of phosphorylated STK32A, adenosine triphosphate (ATP) and magnesium chloride were added to a final concentration of 5 mM in 80 µL of STK32A at 9.5 mg/mL (210 µM). The reaction was monitored by denaturing LC-MS and then the sample was flash frozen. Inline HPLC-SAXS was performed at Diamond beamline b21. The samples were thawed and centrifuged and 45 µL injected onto a Shodex KW-403 column pre-equilibrated in buffer A (25 mM HEPES pH 7.5, 200 mM NaCl, 0.5 mM TCEP, 2.5% glycerol, 1% sucrose), connected to an Agilent 1200 HPLC system. Data was processed and analysed using ScÅtter software [37]. Buffer subtraction was performed separately for a range of stable *R*_*g*_ values across each intensity peak using the SEC flow-through prior to elution of the protein. Radius of gyration (*R*_*g*_) was determined using both Guinier fitting and by analysis of the P(r) distribution. Maximum particle dimension (*d*_max_) and volume of correlation (*V*_c_) were also determined. Molecular mass of the proteins was determined using the *Q*_R_ method [24]. The missing loops from the STK32A crystal structure (PDB 4FR4) were modelled using MODELLER [38] and then FoXS [39] was used to calculate scattering curves from the generated model and compare them to the experimental data. The reduced experimental SAXS data were input into Gasbor [40] and a dummy residue model generated. SUPCOMB [41] was then run to superpose the final models against the crystal structure.

### Inhibitor screening

A library of kinase inhibitors was screened at 10 µM concentration against the purified STK32A protein at 2 µM by differential scanning fluorimetry (DSF), using published procedures [42]. Compounds that significantly increased the melting temperature of the protein were considered hits.

### Cell Lines

SKOv3, HeLa, MCF7 and HEK-293T cell lines were obtained from the American Tissue Type Culture Collection (ATCC). SKOv3 cells were maintained in McCoy’s 5A medium (Invitrogen) while HeLa, MCF7 and HEK-293T were maintained in Dulbecco’s Modified Eagle Medium (DMEM) (Invitrogen). Growth medium was supplemented with 10% (v/v) fetal bovine serum (Invitrogen) and 1% (v/v) penicillin/streptomycin (Invitrogen) and cells were grown at 37**°**C and 5% CO2. SKOv3 cell lines stably expressing STK32A, STK32B, and STK32C were grown in media containing 50 ng/L puromycin (Sigma-aldrich).

### Gateway cloning

Lenti-viral and GFP vectors were generated using the Gateway System (Invitrogen) according to manufacturer’s instructions. STK32A, STK32B and STK32C cDNAs were subcloned into pDONR221 vector (Invitrogen) according to manufacturer’s instruction and then transferred to pLX302 lentiviral destination vector (addgene) or pDEST-CMV-N-EGFP vector (pDEST-CMV-N-EGFP was a gift from Robin Ketteler (Addgene plasmid # 122842) using recombination utilising the LR-Clonase (Invitrogen) as per manufacturer’s instructions. DNA sequences of all cloned cDNAs were verified with direct sequencing.

### Transfections

GFP plasmid transfections were conducted using Fugene HD (Promega) according to the manufacturer’s instructions. Briefly, plasmid DNA and Fugene HD were diluted in Optimem (Invitrogen) in a 1:3 ratio (DNA:Fugene) and incubated for 20 minutes. Cells were transfected and grown at 37°C and 5% CO2 for 48 h or 72 h before they were analysed on the Zeiss Observer Z1 Microscope using a 40x oil objective or a 20x objective. Images were analysed on ImageJ software.

### Lenti-virus mediated stable cell lines generation

Packaging cells (HEK-293T) were co-transfected using Fugene HD (Promega) according to the manufacturer’s instructions with the packaging vector psPAX2, the envelope plasmid pMD2.G (both were gifts from Dr Didier Trono; Addgene plasmids 12260 and 12259) and pLX302 (a gift from Dr David Root; Addgene plasmid 25896) containing the gene of interest. Following 72 h the HEK293-T medium containing the virus was collected, filtered through an 0.45 μM Minisart NML Syringe Filter (Sartorius) and transferred on SKOv3 plated 24 h before infection. The medium was replaced after 72 h and cells containing the integrated virus were selected with Puromycin (50ng/μl) (Sigma).

### Immunofluorescence

Cells were grown to 80% confluence on cover slips in 12-well plates prior to fixation with 0.5ml 4% methanol-free formaldehyde (ThermoFisher Scientific) for 4 minutes at room temperature. Permeabilisation was achieved using 0.5 mL 100% ice-cold ethanol and incubation at −20°C overnight. Ethanol was removed and 1ml wash buffer (1x TBS, 0.2% Triton X100 and 0.04% SDS, Sigma) was added to the wells for 5 minutes. Cells were blocked in 1ml blocking buffer (3% bovine serum albumin diluted in 1xTBS) for 1 hour at room temperature. Cover slips were incubated with 60μl primary antibody (rabbit anti-V5-tag (Abcam) diluted in blocking buffer for 1 hour at room temperature. Three washes were performed with 1ml wash buffer for 5 minutes each. Secondary antibodies (Alexa Fluor 488 donkey anti-rabbit, Invitrogen) were diluted in blocking buffer and 60μl was added to each cover slip for 1 hour at room temperature in the dark. Cells were washed as before prior to mounting the coverslips with mounting medium containing DAPI (Vectashield) on microscope slides (ThermoScientific) sealed with nail varnish at the edges. Slides were stored at 4°C in the dark until analysis. Images were analysed on the Zeiss Observer Z1 Microscope using a 40x oil objective or a 20x objective. Images were analysed on ImageJ software.

## Acknowledgements

The SGC is a registered charity (number 1097737) that receives funds from AbbVie, Bayer Pharma AG, Boehringer Ingelheim, Canada Foundation for Innovation, Eshelman Institute for Innovation, Genome Canada, Innovative Medicines Initiative (EU/EFPIA) [ULTRA-DD grant no. 115766], Janssen, Merck KGaA Darmstadt Germany, MSD, Novartis Pharma AG, Ontario Ministry of Economic Development and Innovation, Pfizer, São Paulo Research Foundation-FAPESP, Takeda, and Wellcome [106169/ZZ14/Z].

## Supplementary Material

**Supplementary Figure 1.**
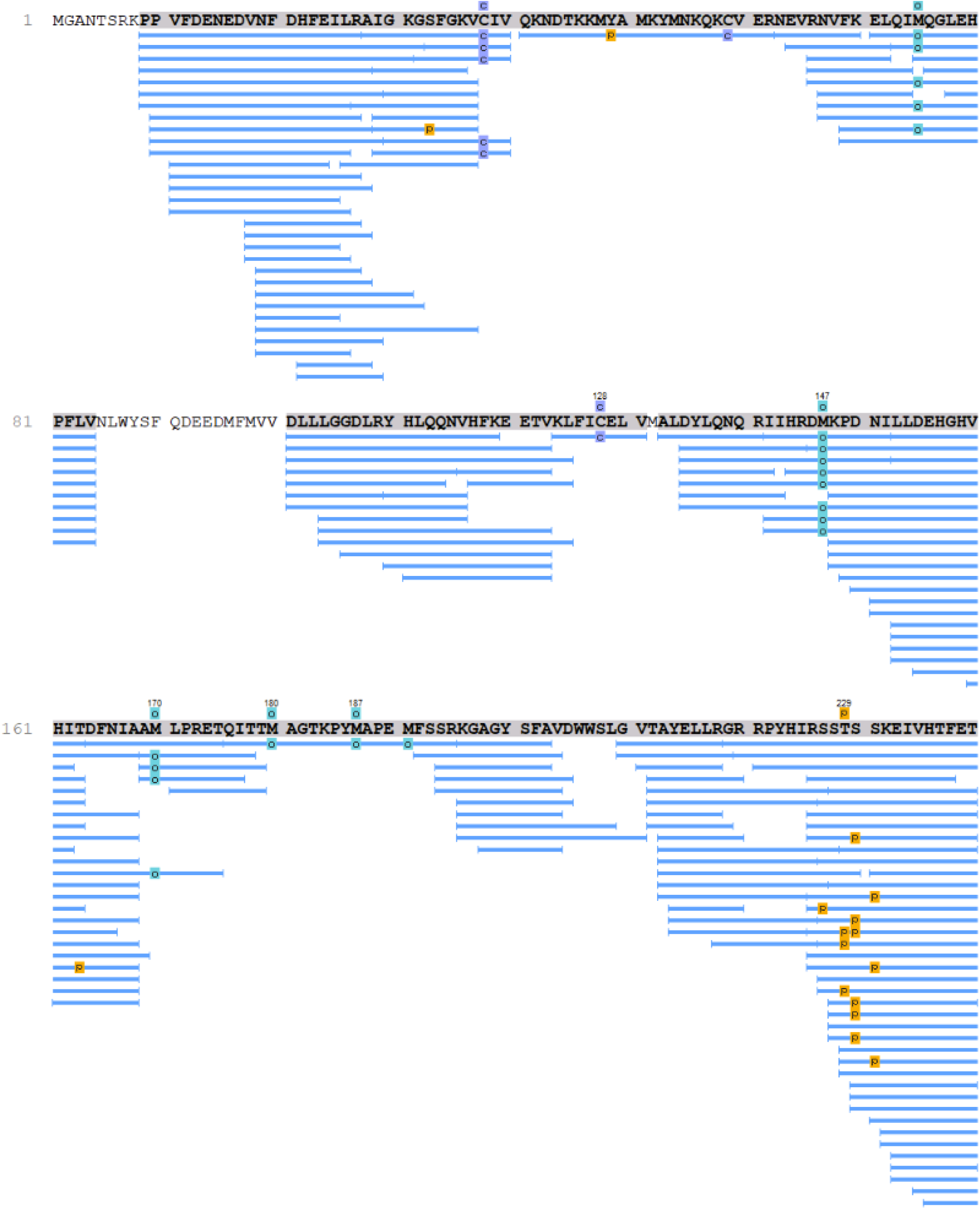
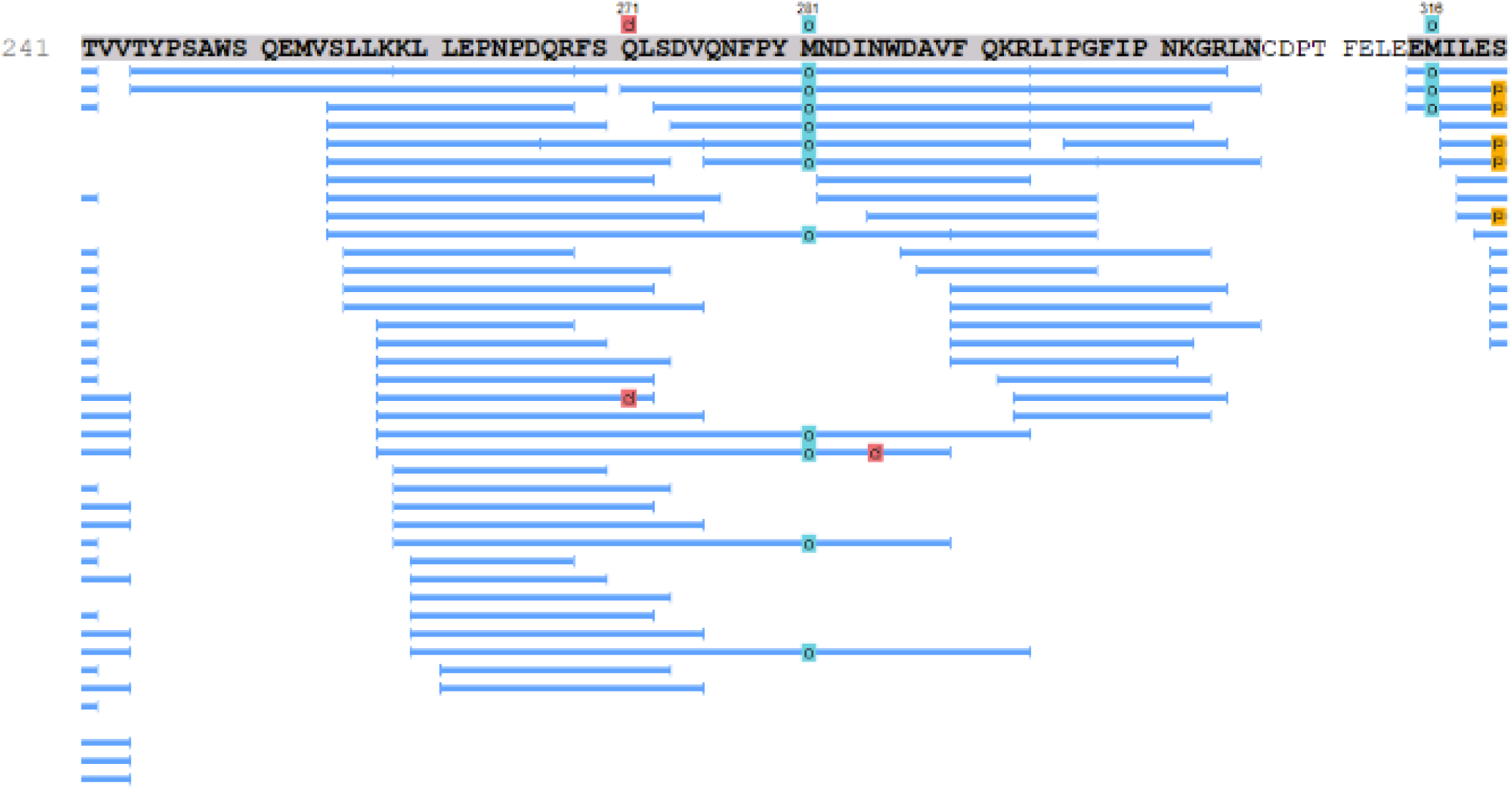
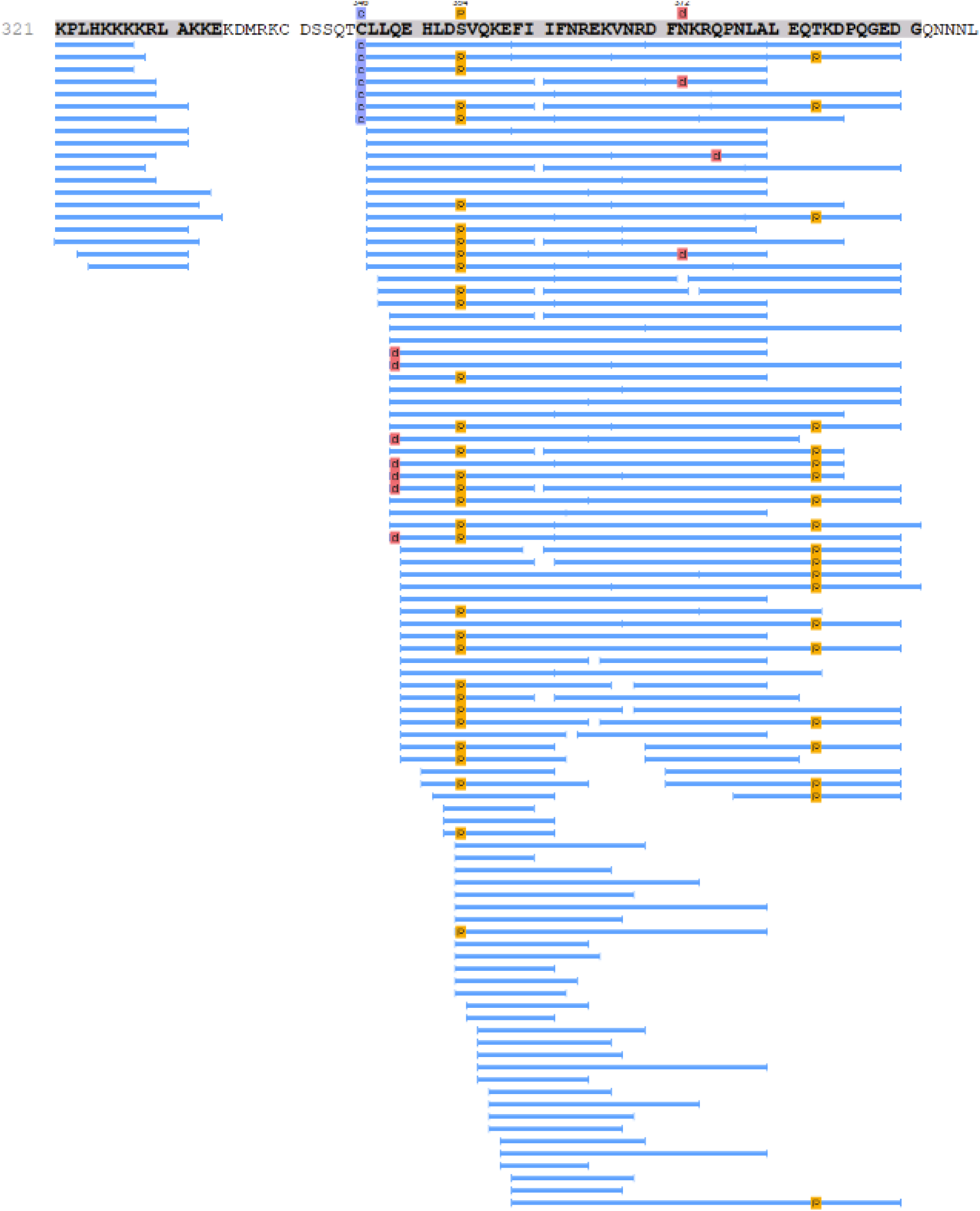
STK32A autophosphorylation peptide coverage. Peptide coverage of STK32A following autophosphorylation and digestion with elastase. Modifications are represented by the following: c (purple) = carbamidomethylation (+57.02 Da), d (red) = deamidation (NQ) (+0.98 Da), o (blue) = oxidation (M) (+15.99 Da) and p (orange) = phosphorylation (STY) (+79.97 Da).

**Supplementary Figure 2.**
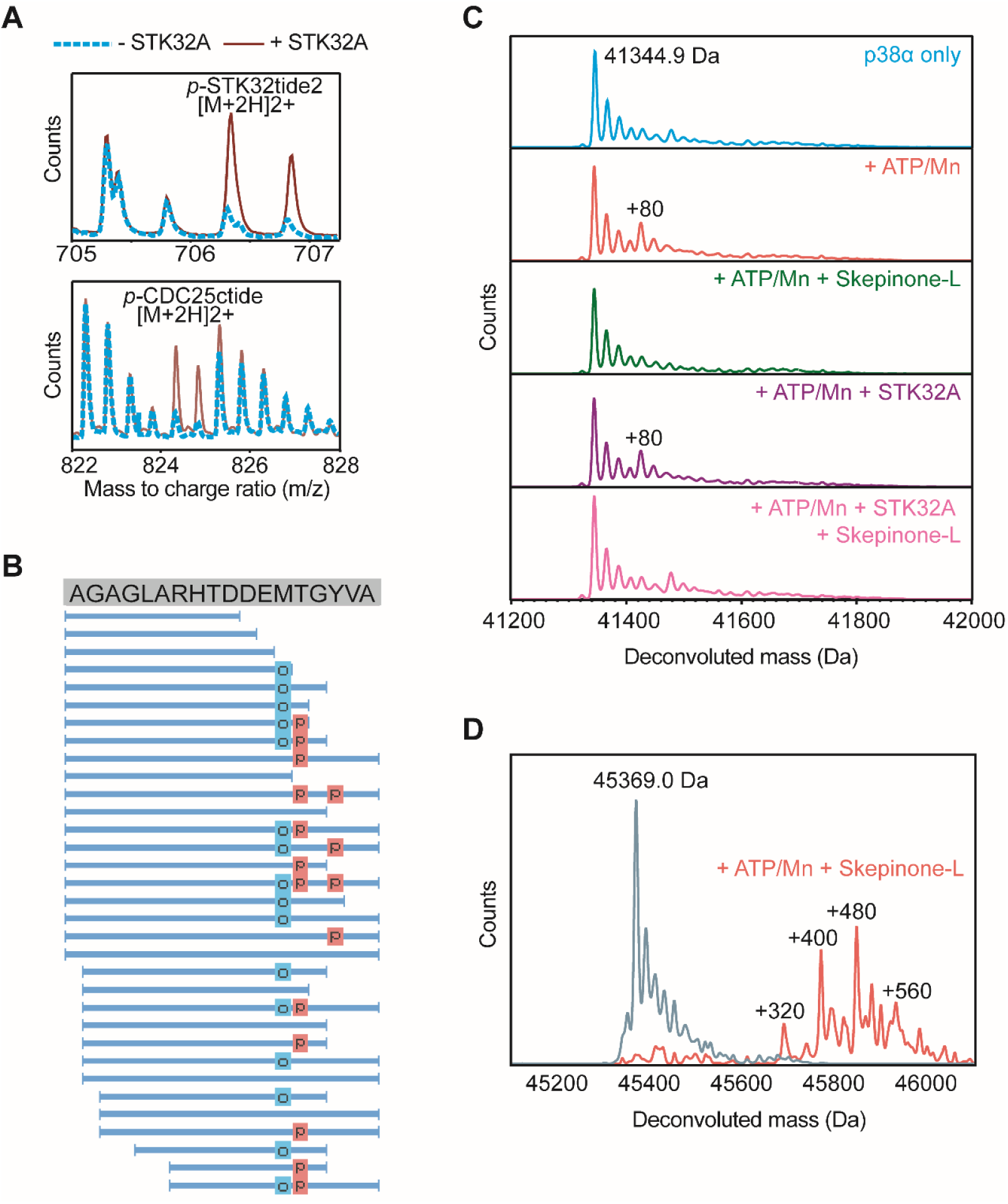
*In vitro* phosphorylation of various substrates by STK32A. **(A)** LC-MS analysis of synthetic consensus peptide substrate STK32tide2 (top) and CDC25ctide (bottom) after phosphorylation by STK32A (red solid line) compared to control without STK32A (blue dotted line). **(B)** Peptide coverage of p38MAPKtide synthetic peptide following phosphorylation by STK32A and digestion using elastase. Blue boxes indicate the position of oxidated residue and red boxes indicate the position of a phosphorylated residue. Supporting peptides are given in Supplementary Table 1. **(C)** Deconvoluted mass spetra of p38α (MAPK14) (top) and following incubation with ATP/Mn +/- STK32A and the specific p38α inhibitor Skepinone-L. **(D)** Deconvoluted mass spectrum of STK32A prior to (grey) and following (red) autophosphorylation in the presence of Skepinone-L.

**Supplementary Figure 3.**
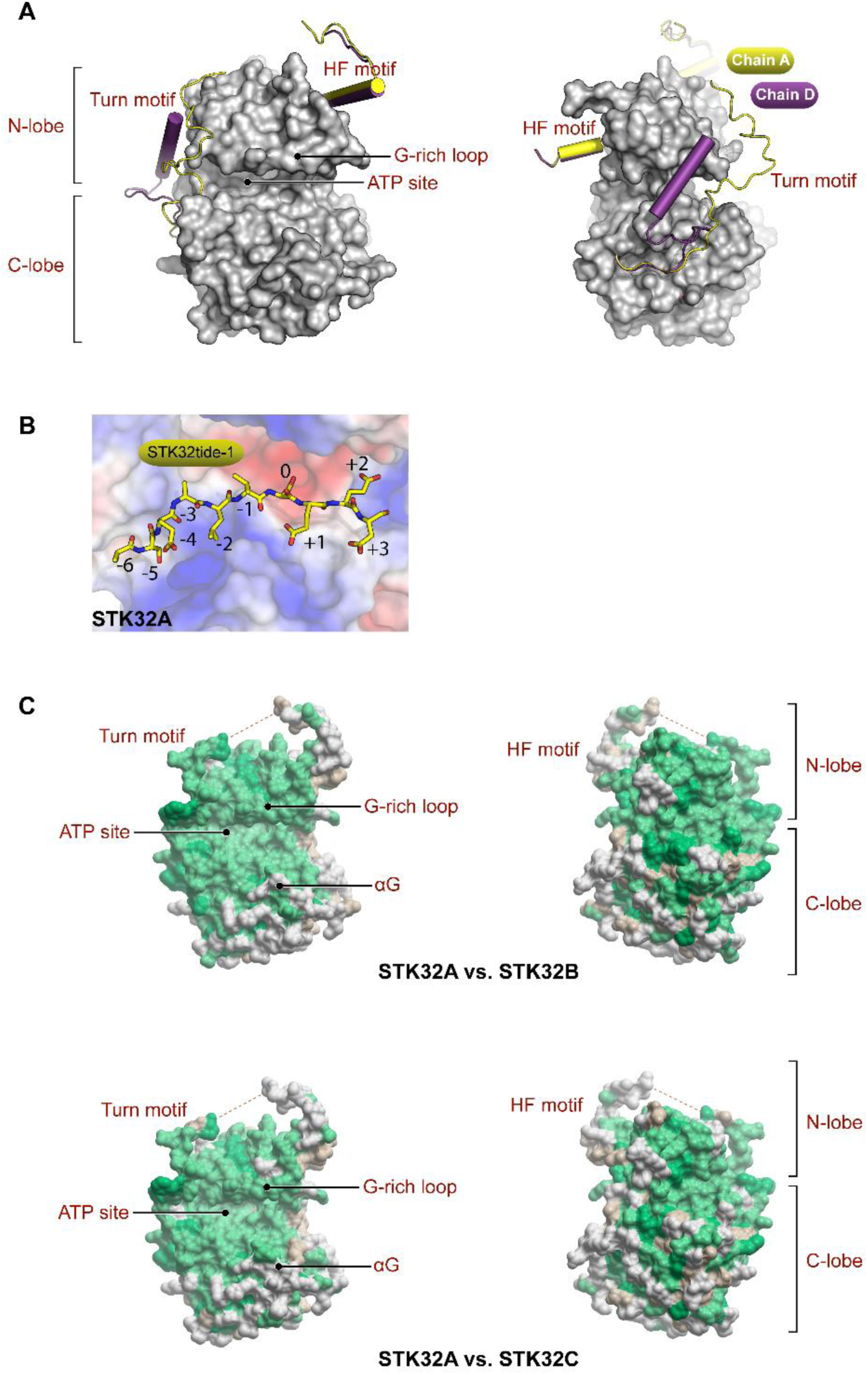
Crystal structure of STK32A. **(A)** Positioning of the turn motif in human STK32A was found in two conformations in the crystal structure (chain A = yellow, chain D = purple). **(B)** Modelling of STK32tide-1 (yellow) in the substrate-binding cleft of STK32A (surface electrostatic potential where blue = positive, red = negative, white = neutral charged). **(C)** Sequence conservation of human STK32B (left) and STK32C (right) mapped onto the crystal structure of STK32A (green = conserved residues, white = non-conserved).

**Supplementary Figure 4.**
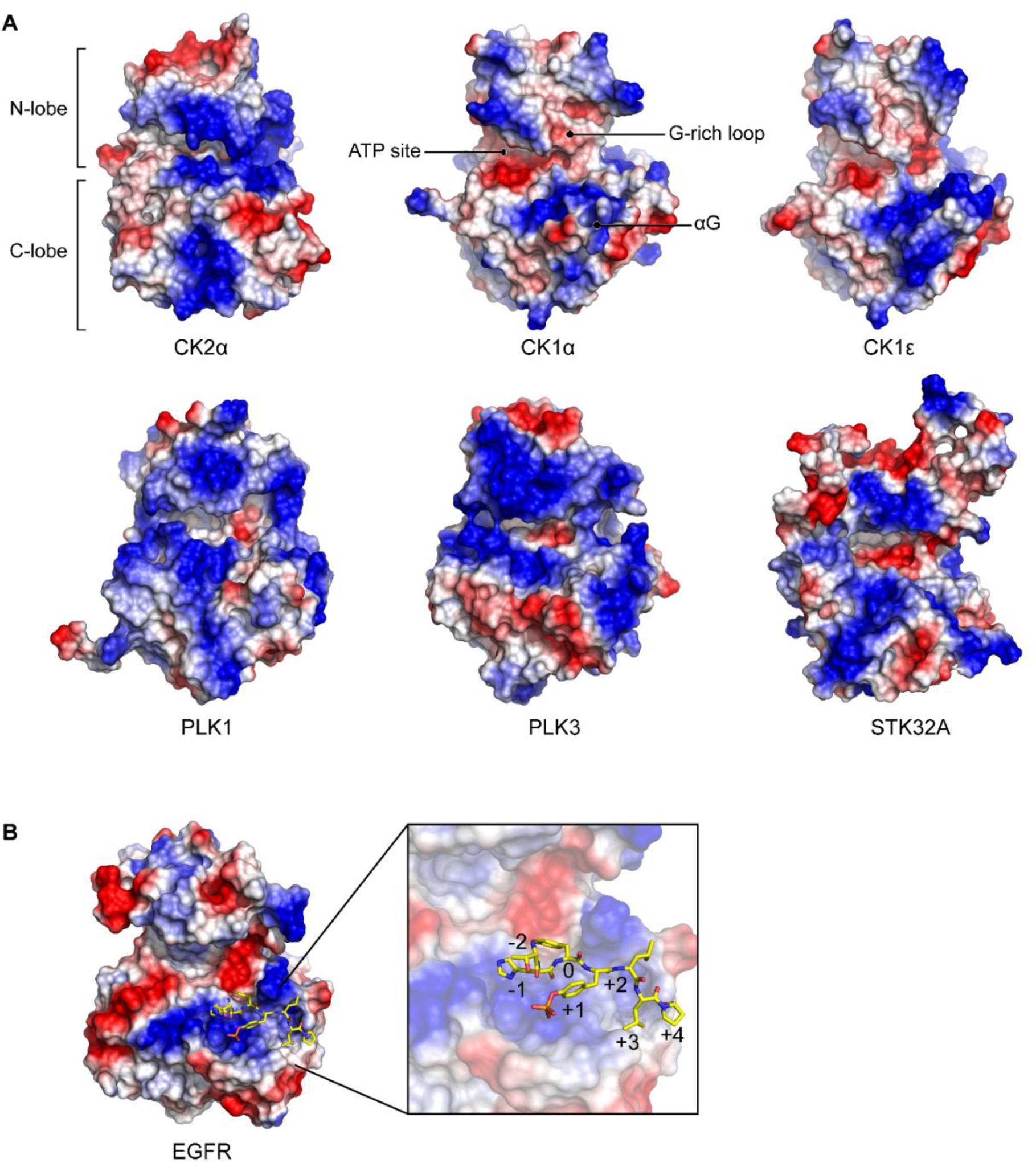
Structural comparison of acidophilic kinases. **(A)** Crystal structures of various known acidophilic serine/threonine kinases coloured by surface charge, where blue indicates positive charge and red indicates negative charge, including casein kinase 2 subunit alpha (CK2α; PDB 1DAW), casein kinase 1 isoform alpha (CK1α; PDB 5IH4), casein kinase 1 isoform epsilon (CK1ε; PDB 4HOK), polo-like kinase 1 (PLK1; PDB 2RKU), polo-like kinase 3 (PLK3; PDB 4B6L) and STK32A (PDB 4FR4). **(B)** Crystal structure of the acidophilic tyrosine kinase epidermal growth factor receptor in complex with a phosphorylated primed substrate (PDB 4R3P), coloured by surface charge as before.

**Supplementary Figure 5.**
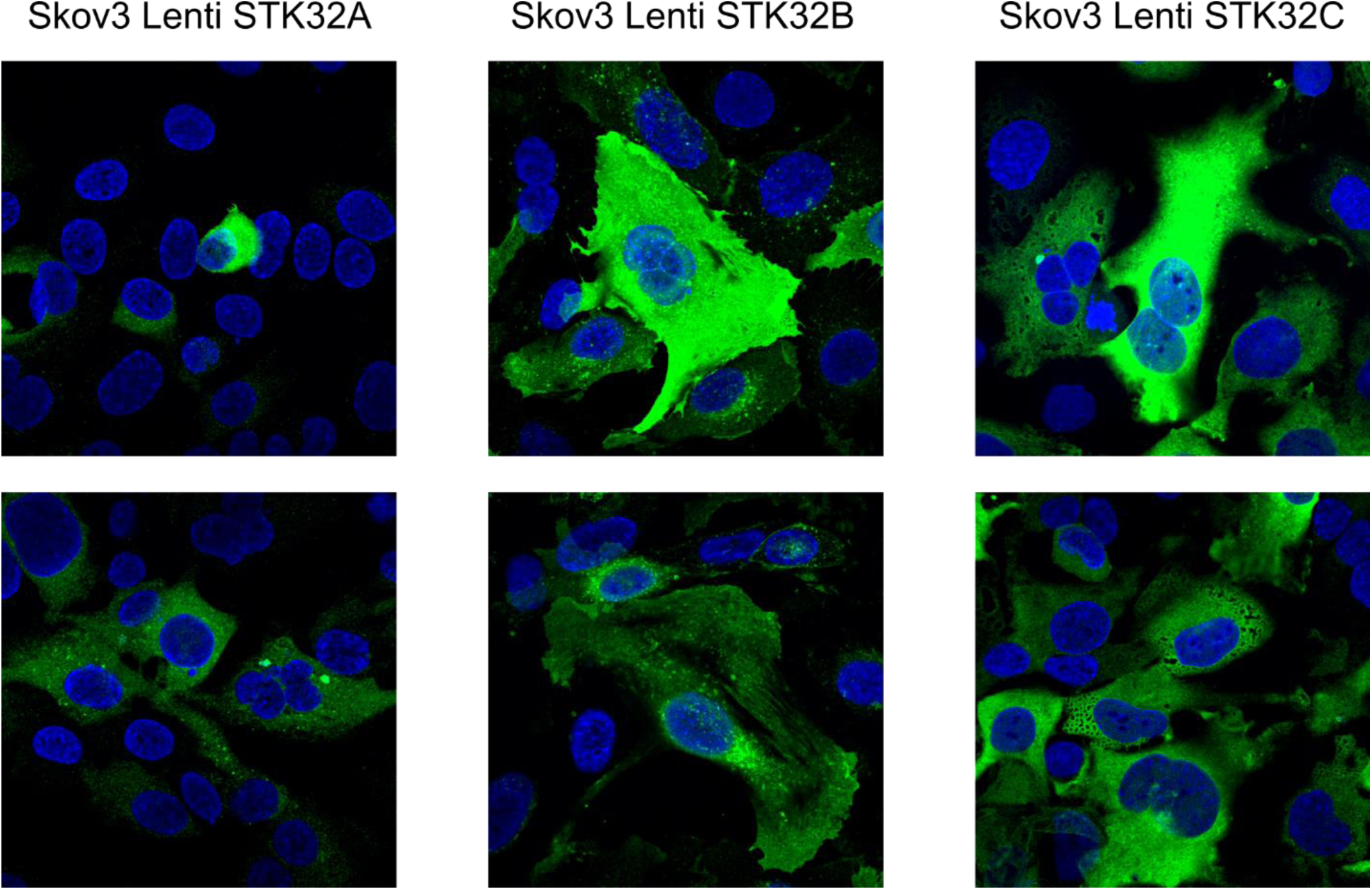
Subcellular localisation of STK32A, STK32B, STK32C. Skov3 cells stably over-expressing V5-tagged STK32A, STK32B or STK32C were fixed and stained for immunofluorescence. Representative images are shown with Hoechst nuclear stain (blue) and anti-V5 antibody (green) showing cytoplasmic but not nuclear localisation.

**Supplementary Figure 6.**
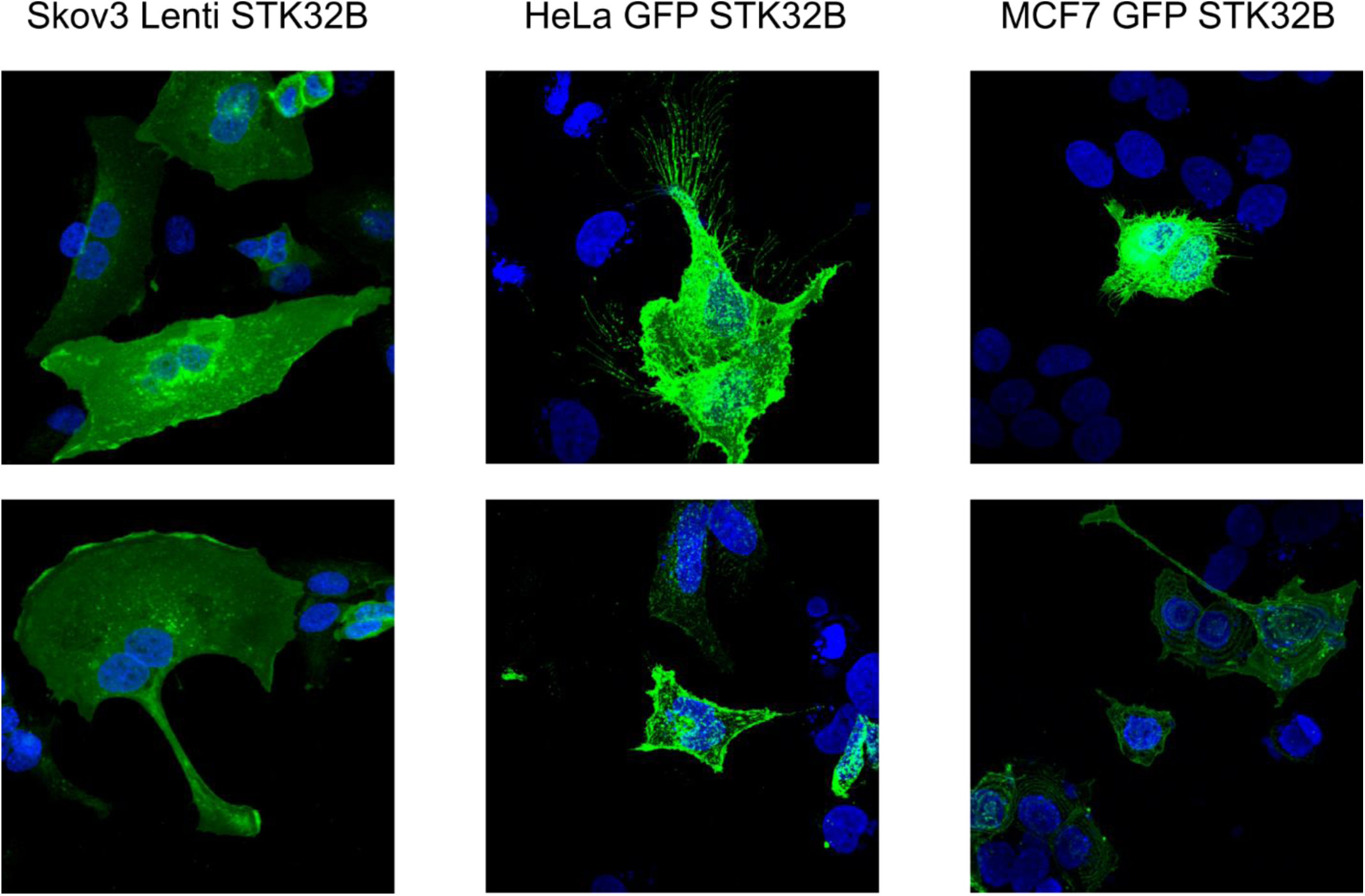
Subcellular localisation of STK32B in different cell lines. Skov3 cells stably over-expressing V5-tagged STK32B or HeLa and MCF7 cells expressing GFP-tagged STK32B were fixed and stained for immunofluorescence. Representative images are shown with Hoechst nuclear stain (blue) and anti-V5 antibody (green) showing cytoplasmic but not nuclear localisation.

**Supplementary Table 1.** p38MAPKtide phosphorylation by STK32A, supporting peptides.

| Peptide | -10lgP | Mass | Length | ppm | m/z | RT | Area<br>Sample 1 | Start | End | PTM |
| --- | --- | --- | --- | --- | --- | --- | --- | --- | --- | --- |
| AGAGLARHTDDEMTGYVA | 43.36 | 1833.8315 | 18 | 7.4 | 917.9264 | 28.39 | 5.92E+05 | 1 | 18 |  |
| AGAGLARHTDDEM(+15.99)T(+79.97)G.Y | 39.09 | 1596.6239 | 15 | 4.4 | 799.3197 | 21.72 | 3.79E+08 | 1 | 15 | Oxidation (M);<br>Phosphorylation<br>(STY) |
| AGAGLARHTDDE.M | 37.6 | 1211.553 | 12 | 4.3 | 606.7841 | 20.8 | 4.21E+06 | 1 | 12 |  |
| A.GAGLARHTDDEMTGYVA | 36.32 | 1762.7944 | 17 | 6.5 | 588.607 | 28.15 | 6.02E+05 | 2 | 18 |  |
| A.GAGLARHTDDEMT(+79.97)G.Y | 36.02 | 1509.5919 | 14 | 3.1 | 755.8027 | 23.83 | 3.32E+06 | 2 | 15 | Phosphorylation<br>(STY) |
| AGAGLARHTDDEMT(+79.97)GY(+79.97)VA | 35.01 | 1993.7642 | 18 | 5.4 | 997.891 | 30.29 | 6.04E+07 | 1 | 18 | Phosphorylation<br>(STY) |
| AGAGLARHTDDEMT(+79.97)G.Y | 34.65 | 1580.629 | 15 | 4.2 | 791.3221 | 23.62 | 2.51E+08 | 1 | 15 | Phosphorylation<br>(STY) |
| AGAGLARHTDDEM(+15.99)TGY.V | 34.64 | 1679.7209 | 16 | 7.5 | 840.8708 | 24.07 | 4.38E+05 | 1 | 16 | Oxidation (M) |
| AGAGLARHTDDEM(+15.99).T | 33.76 | 1358.5885 | 13 | 3.4 | 680.3013 | 20.98 | 7.73E+06 | 1 | 13 | Oxidation (M) |
| AGAGLARHTDDEM(+15.99)TGYVA | 33.72 | 1849.8264 | 18 | 5.1 | 617.6169 | 25.57 | 1.79E+08 | 1 | 18 | Oxidation (M) |
| AGAGLARHTDDEM(+15.99)TG.Y | 33.48 | 1516.6576 | 15 | 3.7 | 759.336 | 21.22 | 1.99E+07 | 1 | 15 | Oxidation (M) |
| AGAGLARHTDDEM(+15.99)T(+79.97)GYVA | 33.11 | 1929.7927 | 18 | 7 | 965.9067 | 26.55 | 7.19E+07 | 1 | 18 | Oxidation (M);<br>Phosphorylation<br>(STY) |
| AGAGLARHTDDEM(+15.99)TGY(+79.97)VA | 32.64 | 1929.7927 | 18 | 8.4 | 965.9081 | 23.49 | 1.73E+07 | 1 | 18 | Oxidation (M);<br>Phosphorylation<br>(STY) |
| AGAGLARHTDD.E | 32.3 | 1082.5105 | 11 | 4.4 | 542.2629 | 20.11 | 3.05E+06 | 1 | 11 |  |
| A.GAGLARHTDDEM(+15.99)TGYVA | 32.28 | 1778.7893 | 17 | 6.3 | 593.9385 | 25.11 | 5.06E+06 | 2 | 18 | Oxidation (M) |
| A.GAGLARHTDDEMTG.Y | 31.99 | 1429.6256 | 14 | 4.4 | 715.8206 | 23.13 | 8.71E+05 | 2 | 15 |  |
| AGAGLARHTDDEM(+15.99)T(+79.97)GY(+79.97)VA | 31.79 | 2009.7592 | 18 | 7.7 | 1005.8907 | 26.8 | 2.72E+07 | 1 | 18 | Oxidation (M);<br>Phosphorylation<br>(STY) |
| AGAGLARHTDDEM(+15.99)T(+79.97).G | 31.39 | 1539.6024 | 14 | 4.6 | 770.8091 | 21.61 | 1.50E+06 | 1 | 14 | Oxidation (M);<br>Phosphorylation<br>(STY) |
| AGAGLARHTDDEMTGY(+79.97)VA | 30.33 | 1913.7979 | 18 | 9.2 | 638.9434 | 25.57 | 2.18E+06 | 1 | 18 | Phosphorylation<br>(STY) |
| AGAGLARHTD.D | 29.29 | 967.4835 | 10 | 1.7 | 484.748 | 20.9 | 6.63E+06 | 1 | 10 |  |
| G.AGLARHTDDEMTGYVA | 27.73 | 1705.7729 | 16 | 4.9 | 569.5989 | 28.02 | 9.22E+04 | 3 | 18 |  |
| AGAGLARHTDDEMTG.Y | 27.68 | 1500.6627 | 15 | 3.6 | 751.3385 | 22.92 | 4.74E+06 | 1 | 15 |  |
| G.AGLARHTDDEM(+15.99)TG.Y | 26.74 | 1388.599 | 13 | 3.7 | 695.3067 | 20.39 | 7.06E+05 | 3 | 15 | Oxidation (M) |
| AGAGLARHTDDEM.T | 26.23 | 1342.5935 | 13 | 3.8 | 448.5385 | 23.67 | 2.21E+06 | 1 | 13 |  |
| AGAGLARHTDDEMT(+79.97)GYVA | 25.77 | 1913.7979 | 18 | 2 | 638.9388 | 26.8 | 9.72E+06 | 1 | 18 | Phosphorylation (STY) |
| A.GAGLARHTDDEM(+15.99)TG.Y | 25.47 | 1445.6205 | 14 | 3.8 | 723.8176 | 20.56 | 1.42E+06 | 2 | 15 | Oxidation (M) |
| G.LARHTDDEM(+15.99)TG.Y | 23.04 | 1260.5404 | 11 | 4 | 631.2776 | 19.05 | 9.12E+04 | 5 | 15 | Oxidation (M) |
| A.RHTDDEMT(+79.97)G.Y | 22.96 | 1140.3907 | 9 | 4.2 | 571.2029 | 20.29 | 3.01E+06 | 7 | 15 | Phosphorylation (STY) |
| A.GAGLARHTDDEMT.G | 22.39 | 1372.6041 | 13 | 4.7 | 458.5424 | 21.27 | 1.32E+07 | 2 | 14 |  |
| AGAGLARHTDDEM(+15.99)T.G | 20.42 | 1459.6361 | 14 | 4.7 | 730.826 | 21.36 | 1.05E+06 | 1 | 14 | Oxidation (M) |
| G.AGLARHTDDEMT(+79.97)GYVA | 19.93 | 1785.7393 | 16 | 7.4 | 596.2559 | 29.46 | 6.90E+04 | 3 | 18 | Phosphorylation (STY) |
| A.GAGLARHTDDEM(+15.99)T(+79.97)GYVA | 19.84 | 1858.7556 | 17 | 9.6 | 620.5961 | 22.87 | 1.47E+06 | 2 | 18 | Oxidation (M); Phosphorylation (STY) |
| A.RHTDDEM(+15.99)T(+79.97)GYVA | 19.59 | 1489.5544 | 12 | 4.7 | 745.7852 | 23.79 | 1.71E+06 | 7 | 18 | Oxidation (M); Phosphorylation (STY) |

**Supplementary Table 2.** Conserved kinase domain residues of AKT2, PKA and STK32A. Conserved residues in the kinase domain core and ATP binding site in active kinases [1] shown for AGC kinases Akt2, PKA and STK32A. Non-conserved residues are highlighted in yellow.

|  | Type | Akt2 | PKA | STK32A |
| --- | --- | --- | --- | --- |
| 1 | h | L158 | L49 | I29 |
| 2 | G | 159 | 50 | 30 |
| 3 | G | 161 | 52 | 32 |
| 4 | V | 166 | 57 | V37 |
| 5 | h | Y178 | Y69 | Y49 |
| 6 | A | 179 | 70 | 50 |
| 7 | h | M180 | M71 | M51 |
| 8 | K | 181 | 72 | 52 |
| 9 | E | 200 | 91 | 71 |
| 10 | L | 204 | 95 | M76 |
| 11 | L | 215 | 106 | 86 |
| 12 | h | F227 | M118 | M98 |
| 13 | V | 228 | 119 | V99 |
| 14 | E | 230 | 121 | D101 |
| 15 | h | A232 | V123 | L103 |
| 16 | L | 237 | M128 | 108 |
| 17 | h | I259 | I150 | L130 |
| 18 | h | L263 | F154 | L134 |
| 19 | H | 267 | 158 | Q138 |
| 20 | h | V271 | L162 | I142 |
| 21 | h | V272 | I163 | I143 |
| 22 | H | Y273 | Y164 | 144 |
| 23 | R | 274 | 165 | 145 |
| 24 | D | 275 | 166 | 146 |
| 25 | I | I276 | L167 | M147 |
| 26 | K | 277 | 168 | 148 |
| 27 | N | 280 | 171 | 151 |
| 28 | h | L281 | L172 | I152 |
| 29 | h | M282 | L173 | L153 |
| 30 | h | L283 | I174 | L154 |
| 31 | I | I289 | I180 | V160 |
| 32 | K | 290 | Q181 | H161 |
| 33 | I | I291 | V182 | 162 |
| 34 | D | 293 | 184 | 164 |
| 35 | F | 294 | 185 | 165 |
| 36 | G | 295 | 186 | N166 |
| 37 | P | 319 | 207 | 189 |
| 38 | E | 320 | 208 | 190 |
| 39 | D | 332 | 220 | 205 |
| 40 | W | 334 | 222 | 207 |
| 41 | G | 337 | 225 | 210 |
| 42 | h | L384 | L272 | L260 |
| 43 | L | 385 | 273 | 261 |
| 44 | R | 392 | 280 | 268 |

**Supplementary Table 3.**
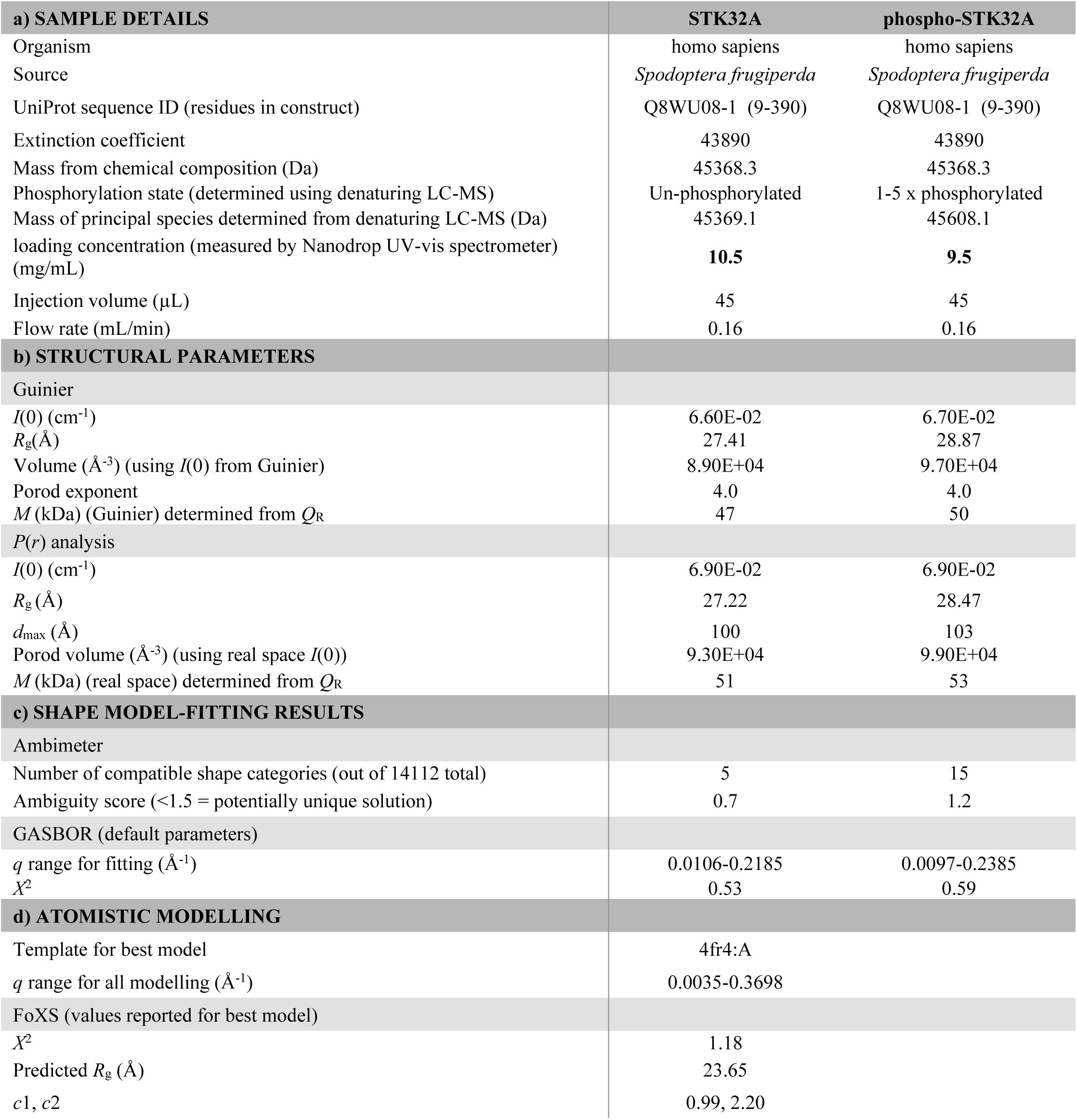
HPLC-SAXS for STK32A.

